# Structural basis for assembly and lipid-mediated gating of LRRC8A:C volume-regulated anion channels

**DOI:** 10.1101/2022.07.31.502239

**Authors:** David M. Kern, Julia Bleier, Somnath Mukherjee, Jennifer M. Hill, Anthony A. Kossiakoff, Ehud Y. Isacoff, Stephen G. Brohawn

## Abstract

Leucine-rich repeat-containing protein 8 (LRRC8) family members form volume regulated anion channels activated by hypoosmotic cell swelling. LRRC8 channels are ubiquitously expressed in vertebrate cells as heteromeric assemblies of LRRC8A (Swell1) and LRRC8B-E subunits. Channels of different subunit composition have distinct properties that explain the functional diversity of LRRC8 currents implicated in a broad range of physiology. However, the basis for heteromeric LRRC8 channel assembly and function is unknown. Here, we leverage a fiducial-tagging strategy to determine single-particle cryo-electron microscopy structures of heterohexameric LRRC8A:C channels in detergent micelles and lipid nanodiscs in three conformations. LRRC8A:C channels show pronounced changes in channel architecture compared to homomeric channels due to heterotypic cytoplasmic LRR interactions that displace LRRs and the LRRC8C subunit away from the conduction axis and poise the channel for activation. The structures and associated functional studies further reveal that lipids embedded in the channel pore block ion conduction in the closed state. Together, our results provide insight into determinants for heteromeric LRRC8 channel assembly, activity, and gating by lipids.

## Introduction

Volume-regulated anion channels formed by LRRC8 (leucine-rich repeat-containing family member 8) proteins open in response to cell swelling and other stimuli to conduct anions and osmolytes across the cell membrane^1-4^. LRRC8 channels are nearly ubiquitously expressed in mammalian cells and play central roles in the cellular response to osmotic stress^5,6^. LRRC8s additionally contribute to a wide range of physiological processes including learning, memory, and response to stroke in the brain through non-vesicular glutamate neurotransmission ^7-9^, immune cell signaling through cyclic dinucleotides^10-12^, insulin and GABA release from pancreatic beta cells^13-16^, spermatogenesis^17^, control of adipocyte and myocyte size^13,18,19^, and vascular function^20^.

LRRC8 channels in cells are predominantly assembled as heterohexamers of LRRC8A (also known as Swell1), which is required for localization to the plasma membrane, and one or more additional paralogs LRRC8B-E^3,4,8,21^. Subunit composition determines channel properties such as substrate selectivity, conductance, inactivation, rectification, and sensitivity to different regulatory inputs. For example, LRRC8A:B heteromers are only permeable to small anions, while LRRC8A:C, LRRC8A:D, and LRRC8A:E heteromers are differentially permeable to a wide range of substrates including neurotransmitters (aspartate, glutamate, GABA, glycine, and ATP), osmolytes (taurine, myo-inositol), cyclic dinucleotides, and small molecule drugs cisplatin and blasticidin S^8,10-12,22-24^. Differences in LRRC8 channel composition likely underlie the wide range of channel properties observed in different cells^1,2,6^. The mechanistic basis of these differences has remained unknown due to the lack of structural information about heteromeric LRRC8 channel assembly; efforts to date have been limited by pseudosymmetry between subunits and potential variations in stoichiometry that prevent accurate alignment in cryo-EM reconstructions^25^.

Homomeric LRRC8A and LRRC8D channel structures have provided insight into channel architecture, anion selectivity, and small molecule inhibition^25-30^, but mechanisms of LRRC8 channel activation and gating remain unclear. Several lines of evidence suggest separation of cytoplasmic leucine-rich repeat (LRR) domains promotes opening of the channel pore. LRR truncation results in nonfunctional channels^17^, C-terminal fusions increase LRRC8 activity^6,31,32^, Förster resonance energy transfer (FRET) imaging has associated channel activation with LRR separation^33^, structures of LRRC8A homohexamers bound to activating nanobodies display increased LRRC8A LRR separation^30^, and different LRRC8A conformations show correlated expansion from LRR to transmembrane regions^28^. The physical basis of channel gating is unclear because structures of homomeric LRRC8A and LRRC8D channels display apparently open conduction pathways despite being determined under conditions expected to promote closed channels. Structurally unresolved N-termini of LRRC8A may gate the channel^34^, but analogous LRRC8D N-termini that are resolved do not obstruct conduction^29^. Additionally, the physiological relevance of homomeric LRRC8 channels is uncertain; they are insufficient for native channel activity, LRRC8D homomers are nonfunctional, and the LRRC8A homohexamer displays small, atypical currents with altered sensitivity to swelling, ionic strength, and inhibitors^35,36^.

Here, we present cryo-EM structures of heteromeric LRRC8A:C channels in detergent micelles and lipid nanodiscs. These structures and associated functional studies provide a basis for understanding the assembly, function, and gating of physiologically relevant volume-regulated anion channels.

## Results

### Structure determination

We used a fiducial-tagging strategy to overcome the pseudosymmetry of LRRC8A and LRRC8C subunits and facilitate accurate subunit alignment in cryo-EM reconstructions (Supplementary Fig. 1). We selected the BRIL domain (b_562_RIL; an engineered variant of apocytochrome b_562_a) as a fiducial^37-39^ and the first extracellular loop of the LRRC8A subunit as the site of insertion. The loop is disordered in homohexameric LRRC8A structures^28^ and precedes functionally important regions^40^ suggesting BRIL insertion would not compromise channel function. To test this, we compared channel activity in LRRC8 knockout HELA cells (LRRC8A/B/C/D/E -/-) co-expressing either LRRCA and LRRC8C or LRRC8A_BRIL_ and LRRC8C subunits. LRRCA:C and LRRC8A_BRIL_:C channels showed indistinguishable activity, with low basal activity in isotonic solution and swelling-induced currents in hypotonic solution, consistent with functional integrity of LRRC8A_BRIL_-containing channels (Supplementary Fig. 1).

For structural studies, we expressed LRRC8A_BRIL_:LRRC8C channels in insect cells that lack endogenous LRRC8 subunits. LRRC8A_BRIL_-sfGFP and LRRC8C-mCherry proteins were expressed using a single engineered baculovirus. The fluorescent protein tags were fused to LRRC8 C-termini through orthogonal protease cleavage sites to allow for specific tag cleavage. Heteromeric channels were isolated from homomeric channels through sequential anti-mCherry and anti-GFP affinity purification (Supplementary Fig. 2). Purified channels in detergent or reconstituted in lipid nanodiscs were complexed with anti-BRIL antibody fragments (Fabs)^39^ and anti-Fab nanobodies^41,42^ to further increase the mass of the BRIL fiducial marks on LRRC8A subunits for structure determination by cryo-EM.

Single particle reconstructions of LRRC8A_BRIL_:C channels solubilized in glyco-diosgenin (GDN) detergent without applied symmetry resulted in maps in different states with an overall resolution of 3.1-3.2 Å (Fig.1, Supplementary Fig. 3-5, Table 1). Through variability analysis, we identified two conformations with prominent differences in the organization of cytoplasmic leucine rich repeat (LRR) domains. For both conformations, focused local refinements of LRRs or the linker, transmembrane, and extracellular region (“top” of the channel) improved resolution and map features and were used for modeling. Partial models from focused maps were merged following refinement, docked into global refinement maps, and subjected to B factor refinement to generate full channel models. Most of the channel is modeled in each conformation apart from one (or three) LRRC8A LRR domains, N-terminal regions, extracellular loop 1 (including the BRIL-Fab-Nb fiducial), and cytoplasmic loop between linker helix 2 and 3.

**Figure 1.**
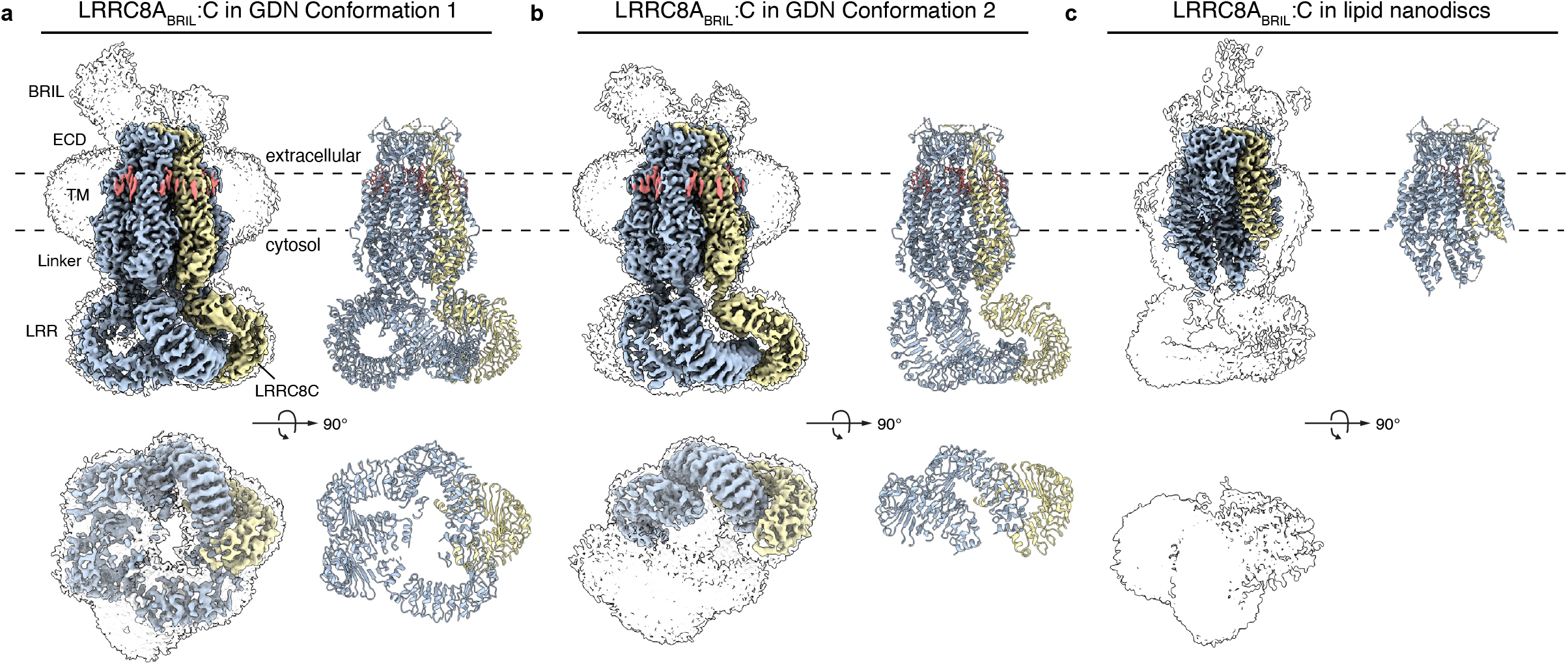
Structures of heterohexameric LRRC8A:C channels. **(a-c)** Cryo-EM density maps (left) and models (right) of LRRC8A:C in **(a**,**b)** detergent micelles and **(c)** lipid nanodiscs viewed from the membrane plane (above) and cytoplasm (below). LRRC8A subunits are blue, LRRC8C subunits are yellow, and lipids are orange.

**Table 1.**
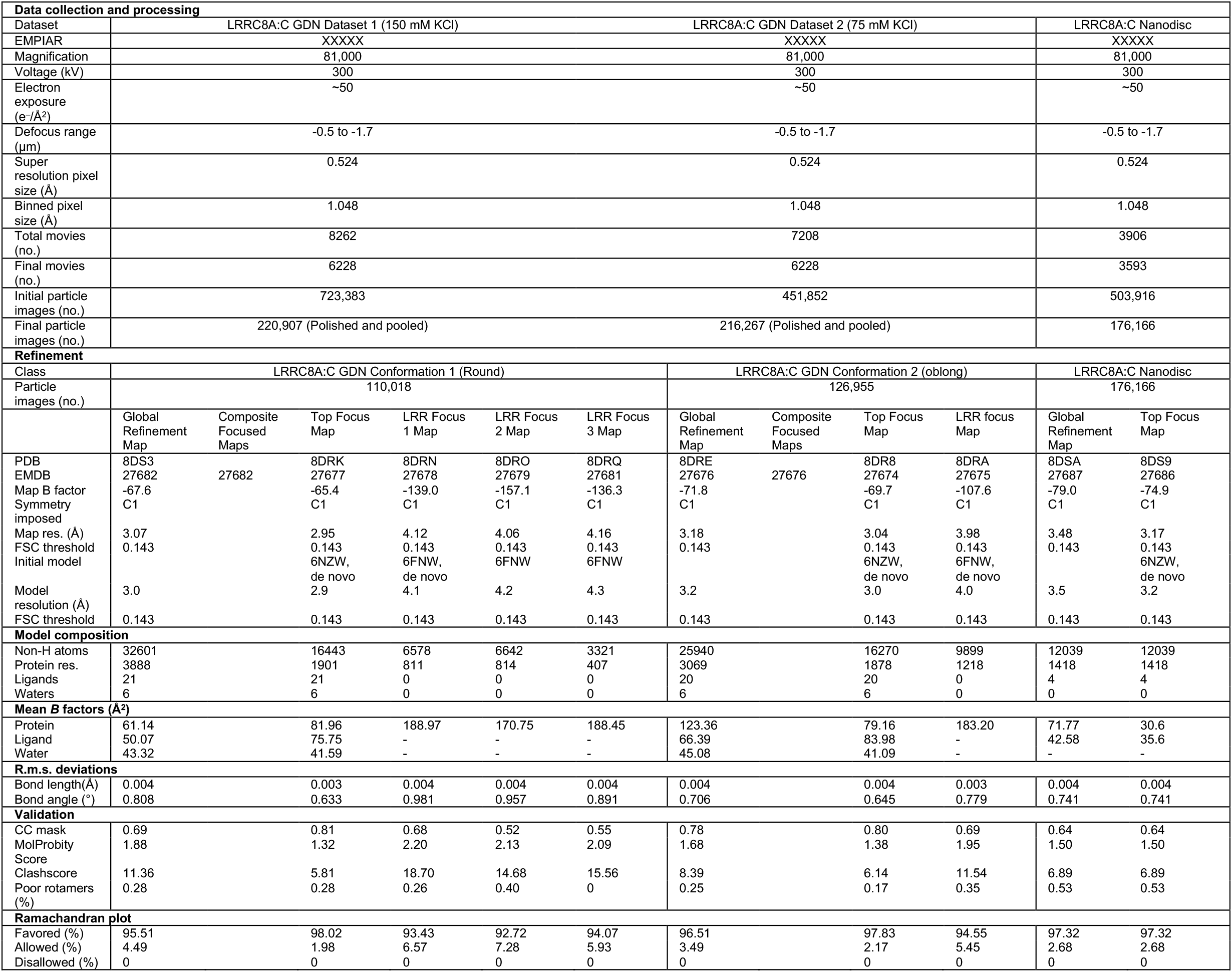
Cryo-EM data collection, refinement, and validation statistics.

A single particle reconstruction of LRRC8A_BRIL_:C channels reconstituted in lipid nanodiscs resulted in a map with an overall resolution of 3.5 Å. Top-focused local refinement led to a 3.2 Å resolution map with improved features that was used for modeling (Fig.1, Supplementary Fig. 6, Table 1). LRRC8A_BRIL_:C exhibits substantially more heterogeneity in the nanodisc sample than in detergent; four linker regions and the six LRR domains are unresolved in the final nanodisc model. We focus our discussion below on the better resolved GDN-solubilized structures unless otherwise noted.

### Overall structure

Final cryo-EM maps and 2D class averages show extra protein density consistent with BRIL-Fab-Nb fiducials above the extracellular region of five LRRC8 subunits (Supplementary Fig. 7a,b). A 5:1 LRRC8A:LRRC8C stoichiometry was confirmed in each structure by differences in backbone organization (amino acids 48-51, 57-95) and density corresponding to amino acid positions that differ between LRRC8A and LRRC8C in the best-ordered regions of the map (Supplementary Fig. 7c). The LRRC8A_BRIL_:C structure demonstrates that modest flexibility between target and engineered fiducial is not necessarily an impediment to accurate subunit alignment.

The LRRC8C subunit consists of extracellular and transmembrane regions, a linker, and a 17-repeat LRR domain (Fig. 2a). The overall fold between LRRC8C and LRRC8A subunits is similar (whole chain Cα r.m.s.d. = 3.5-5.3 Å) and the extracellular domains are nearly invariant between all subunits and structures (r.m.s.d. < 1 Å), suggesting it forms a rigid core of the channel. Comparing LRRC8A:C and LRRC8A channels aligned by their extracellular region reveals LRRC8C subunit incorporation has three major structural consequences with implications for understanding channel function.

**Figure 2.**
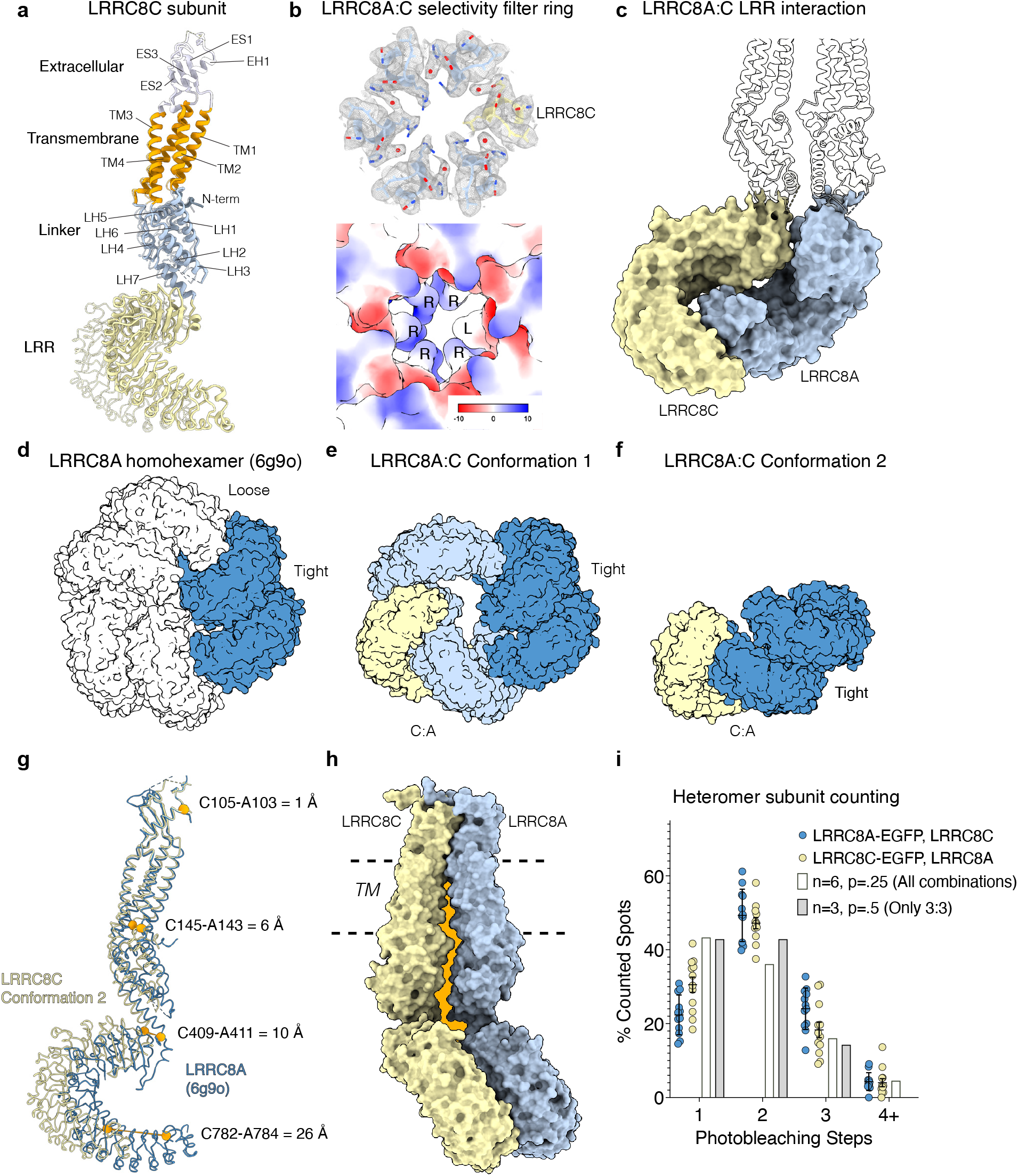
LRRC8C subunit structure and impact on channel architecture. **(a)** LRRC8C subunit structure. Extracellular, transmembrane, linker, and LRR regions are colored white, orange, blue, and yellow, respectively. Secondary structure elements are indicated. Extracellular regions of LRRC8C from conformations 1 and 2 are aligned by their extracellular regions and overlaid with conformation 2 shown as a thin cartoon. **(b)** LRRC8A:C extracellular selectivity filter shown from above with cryo-EM map and model (above, LRRC8C is yellow) and as an electrostatic surface (below). From conformation 1. **(c)** A:C LRR interaction from conformation 1. **(d-f)** Cytoplasmic view of LRR interactions in **(d)** homohexameric LRRC8A **(e)** LRRC8A:C conformation 1, and **(f)** LRRC8A:C conformation 2. LRRC8A LRR pairs interacting through tight interfaces are shown in dark blue, LRRC8C-interacting LRRC8A LRRs in light blue, and LRRC8C LRRs in yellow. **(g)** Superposition of LRRC8A (PDB: 6g9o) and LRRC8C conformation 2 subunits aligned by their extracellular regions showing displacement of LRRC8C from the conduction axis. Distances between residues in the extracellular, transmembrane, linker, and LRR regions are indicated. **(h)** Intersubunit gap (orange) between LRRC8C and LRRC8A subunits in the heteromeric channel. **(i)** Single molecule subunit counting of LRRC8A:C channels. Distribution of photobleaching steps from heteromeric LRRC8A:C channels with EGFP-tagged LRRC8A or LRRC8C expressed in *LRRC8 -/-* HEK293T cells (mean ± sem for n=12 movies with ≥ 50 particles each). Predicted distributions are shown assuming 50% mature EGFP fluorophores and either 3:3 or all possible LRRC8A:C stoichiometries within a hexameric channel.

First, LRRC8C incorporation in heteromeric channels alters the size and chemistry of the extracellular selectivity filter ring (Fig. 2b). In homomeric LRRC8A channels, R103 side chains project from the outer end of the extracellular helix towards the conduction axis to form an electropositive constriction above the pore that partially determines anion selectivity^25^ and is the location for inhibition by DCPIB and related blockers^16,28^. Homomeric LRRC8D displays F143 from the analogous position, resulting in an expanded and hydrophobic filter ring. LRRC8C subunits have a leucine in this position (L105), and its inclusion in the heteromeric LRRC8A:C ring results in a slightly expanded and less electropositive filter ring (Fig. 2b) that is likely to contribute to different substrate permeability and small molecule inhibition of heteromeric channels.

Second, LRRC8C incorporation results in unique interactions between LRRs that alter the arrangement of the entire cytoplasmic region (Fig. 2c). Unlike homotypic LRRC8A or LRRC8D LRR interactions, heterotypic LRRC8A:C interactions involve insertion of an LRRC8A LRR into the concave interior of an LRRC8C LRR. The interaction is similar in both GDN conformations (r.m.s.d. LRRC8A:LRRC8C LRR pair = 1.53 Å), encompasses ∼700-750 Å_2_ of surface area, and is predominantly formed by residues in repeat 1 and 12-17 of LRRC8C and 14-17 of LRRC8A (Fig. 2c).

In homomeric channels, pairs of LRRC8A LRRs form “tight” interactions between their long edges (∼850 Å^2^ buried surface area) and these pairs associate through “loose” interactions (∼300 Å^2^ buried surface area) that abut C-terminal ends and the open long edge of neighboring pairs (Fig. 2d)^25-27^. This results in a trimer of A LRR dimers stabilized by each subunit interacting with each of its immediate neighbors. In heteromeric channels, the A:C LRR interaction is observed either in the context of a single LRRC8A LRR (conformation 1) or dimeric, tight interfaced, pair of LRRC8A LRRs (conformation 2). The different possible arrangements explain the presence of the two conformations in the dataset and the differences in the number of disordered LRRs between the structures. The A:C LRR interaction generates a gap that can only accommodate one additional A:A dimer pair (Fig. 2e,f). In conformation 1, it is separated from the A:C pair by a modified loose A:A interface (similarly sized at ∼350 Å^2^, but with repeats 7-10 instead of 10-13 of the A LRR edge) and a second C:A interface (∼350 Å^2^ buried surface area) (Fig. 2d). This results in 5 stable LRR positions, but insufficient space for the final A LRR to interact with either neighbor. In conformation 2, an A:A pair could fit in either of two open positions, but neither is compatible with formation of an additional A:C or A:A interface (Fig. 2e). This results in 3 stable LRR positions and three flexible A LRRs.

Third, LRRC8C incorporation and formation of the A:C LRR interactions described above result in displacement of the entire C subunit away from the conduction axis and opening of an A:C intersubunit gap (Fig. 2g,h). Cradling an A LRR in its inner surface requires the C LRR to swing outwards 16 to 26 Å, with a larger displacement required to accommodate the A:A pair in conformation 2. This is enabled by a hinge movement about the extracellular region that also displaces the transmembrane and linker region ∼4-6 Å and ∼9-11 Å at their bases, respectively (Fig. 2g). As a result of C subunit displacement, a wide gap is opened between neighboring C and A subunits that extends from the base of the linker to the top of the transmembrane helices (Fig. 2h). This contrasts with other A:A and A:C subunit interfaces that are sealed by protein. The gap is large enough (narrowing to ∼3Å) to permit cytoplasmic ion flux at the linker regions and lipid access between the transmembrane helices.

### LRRC8A:C channel stoichiometry

We observe a 5:1 A:C subunit stoichiometry in heteromeric LRRC8A_BRIL_:C channel structures and did not see evidence of other stoichiometries during particle classification, suggesting 5:1 is the predominant form produced in insect cells under our expression conditions. LRRC8C alone is present in the anti-GFP affinity purification flowthrough, which indicates homomeric LRRC8C channels are produced and LRRC8C subunits may not be limiting for formation of other stoichiometries. Still, we do not identify any obvious structural impediment to formation of LRRC8A:C channels with alternate stoichiometries.

We therefore analyzed LRRC8A:C stoichiometry in LRRC8 knockout (*LRRC8A/B/C/D/E -/-*) HEK293 cells^8^ by single molecule pulldown of transfected channels and stepwise subunit counting by fluorescent photobleaching (Fig. 2i, Supplementary Fig. 8). We first validated the strategy using homodimeric HVCN1 channels and homohexameric LRRC8A channels as controls. Both were expressed as C-terminal EGFP-HA fusions, extracted with detergent, isolated and immobilized on passivated coverslips using anti-HA affinity capture, and imaged with total internal fluorescence microscopy to observe fluorophore bleaching steps. As expected, the majority of HVCN1 molecules display two bleaching steps and count distributions (Supplementary Fig. 8a) that closely match previously published distributions^43^. Homomeric LRRC8A molecules show a higher average number of bleaching steps and a count distribution consistent with expected homohexameric assembly assuming ∼50% of EGFP molecules are dark due to incomplete fluorophore maturation (Supplementary Fig. 8b).

We next co-expressed LRRC8A and LRRC8C subunits as C-terminal fusions to EGFP-HA or mCherry through orthogonal protease cleavage sites. To isolate heteromeric from potential homomeric species and free fluorophores, channels were extracted in detergent and subjected to a two-step affinity capture and display procedure. Count distributions are similar regardless of whether LRRC8A or LRRC8C was EGFP-tagged and most molecules counted show two or more bleaching steps, demonstrating stoichiometries other than 5:1 LRRC8A:C can be generated in HEK293 cells (Fig. 2i). The data are similarly well fit by calculated expected count distributions for a single 3:3 LRRC8A:C stoichiometry or a mixture of all possible LRRC8A:C stoichiometries (assuming 50% of EGFP molecules are fluorescent based on experiments with homomeric LRRC8A channels (Supplementary Fig. 8b)). We are therefore unable to distinguish between these possibilities based on fit to expected distributions alone. Still, we conclude LRRC8A:C channels can assemble with different stoichiometries under different conditions, consistent with previous subunit counting experiments of LRRC8A:E channels in *Xenopus* oocytes^31^.

### Lipid block of conduction in the LRRC8 pore

Cryo-EM maps of LRRC8A:C display striking lipid-shaped density inside the channel pore at the level of the outer leaflet of the lipid bilayer (Fig. 3a-c). Based on shape and chemical environment, we modeled three phospholipids into these densities with their head groups extending into the hydrophilic extracellular pocket between the selectivity filter ring and transmembrane helices, glycerol backbones at the level of the extracellular ends of TM1, and acyl chains extending down against the hydrophobic surfaces of TM1s from adjacent subunits. Strikingly, the three lipids within the LRRC8A:C pore form a complete seal to ion flux through the channel. A constriction between glycerol backbones narrows the pore to a radius of 1.2∼1.5 Å, smaller than dehydrated Cl^-^ (1.8 Å radius) and other permeant species. Without bound lipids, there is no visible impediment to ion conduction; the narrowest point at the selectivity filter ring measures ∼2.1 Å in radius.

**Figure 3.**
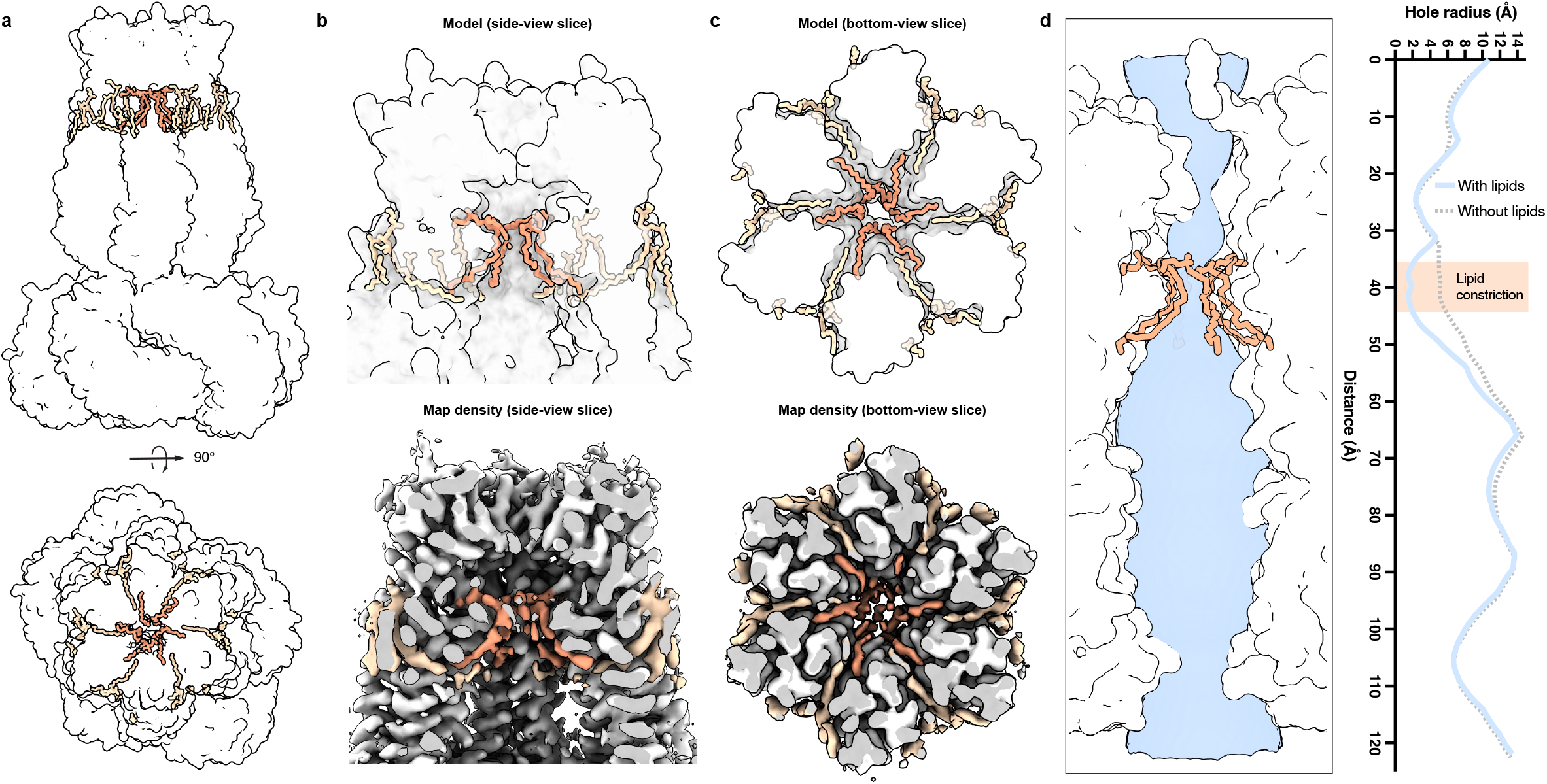
Lipids block conduction in closed LRRC8A:C channels. **(a)** Lipid positions observed in LRRC8A:C structure (using conformation 1) viewed from the membrane (above) and extracellular side (below). Channel is shown as a white surface, external lipids are tan, and internal lipids are orange. **(b**,**c)** Zoomed in **(b)** side view and **(c)** bottom view with cryo-EM map below. **(d)** Channel pore (blue surface, left) and radius (right) calculated with (blue) and without (dotted) bound lipids.

An alternate arrangement in which each lipid is rotated 60º fits the maps less well but is chemically similar, and may be occupied in a subset of particles. Weaker density for lipids compared to adjacent protein may be due to occupancy of this alternate conformations and lipid flexibility within the pore. Density is also present for outer leaflet lipids on the exterior of the channel against intersubunit interfaces, with one phospholipid per interface projecting an acyl chain through intersubunit gaps towards the pore. In the nanodisc structure, lipids are observed in similar positions inside and outside the pore against the best resolved subunits, but not against less well resolved subunits consistent with overall lower resolution due to conformational heterogeneity. Similar headgroup and acyl chain position, relative to the conduction axis, for lipids inside and outside the pore gives the impression of a nearly continuous bilayer through the channel, with pore lipids fenced off from bulk membrane by loose intersubunit protein packing in the upper leaflet.

We hypothesized that if the pore lipids contribute to block of conduction in a closed channel state, mutation of lipid-interacting residues to disfavor lipid association would increase basal activity in isotonic conditions and correspondingly reduce activation by hypotonic swelling (Fig. 4). Recording from basally activated channels is technically challenging, so as a positive control we first recorded LRRC8 channels known to be activated by C-terminal fusion to fluorescent proteins ^6,31-33^. Consistent with activation by C-terminal fusions, LRRC8A-GFP:C-mCherry channels showed nearly ∼20-fold larger basal currents in isotonic solution and ∼20-fold smaller fold-activation in hypotonic solutions compared to wild-type LRRC8A:C channels (Fig. 4b,c and Supplementary Fig. 9).

**Figure 4.**
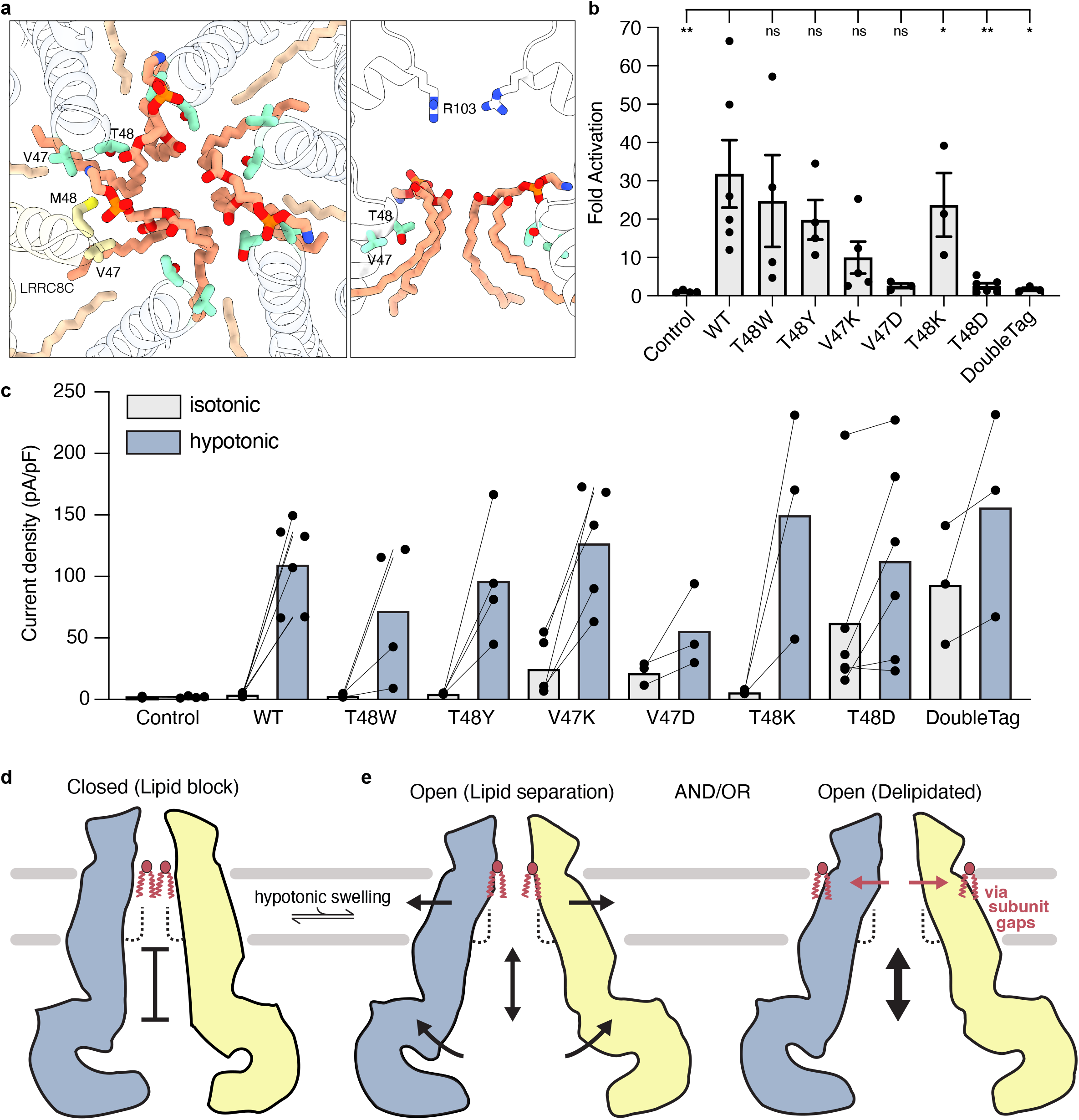
Disrupting lipid interactions activates LRRC8A:C channels. **(a)** Top down and side views of lipid-interacting regions of the channel at the top of TM1 (using conformation 1). Residues mutated in this study, V47 and T48, are indicated. The corresponding LRRC8C residues V47 and M48 as well as the R103 of the selectivity filter are also shown for positional context. **(b)** Fold activation of LRRC8A mutant LRRC8A:C channels by hypotonic cell swelling (I60mV, hypotonic / I60mV, isotonic). Values are mean ± sem for n= 6 (WT, T48D), 5 (V47K), 4 (Control, T48W, T48T), 3 (V47D, T48K, LRRC8A-GFP:C-mCherry). Differences assessed with one-way analysis of variance (ANOVA) with Dunnett correction for multiple comparisons (*P, **P). Note: The Control and WT data are also used for analysis in Supplementary Figure 1. **(c)** Current density at 60 mV in isotonic (gray) and hypotonic (solution). Lines connect data from a single cell. **(d-e)** Model for lipid gating of LRRC8 channels showing two opposing subunits and pore lipids. In a closed resting state **(d)** lipids from the upper leaflet infiltrate the pore and form a seal to ion conduction. LRRC8A:C structures reported here are captured in the closed state. Opening may involve channel expansion to open a path between lipids for ion passage **(e, left)** and/or retreat of lipids from the pore through intersubunit openings **(e, right)**.

We next tested activity of LRRC8A:C channels with mutations in positions that contact pore lipid acyl chains and the bottom of glycerol backbones. Introducing bulky hydrophobic residues in either LRRC8A(T48W):C and LRRC8A(T48Y):C mutant channels resulted in wild-type-like activity, with low basal activity and strong activation by hypotonic swelling (Fig. 4b,c and Supplementary Fig. 9). These data support a model in which hydrophobic interactions promote lipid binding in the pore to block conduction under resting conditions. It further demonstrates bulky residues can be accommodated in this region of LRRC8A. A lysine at this site in LRRC8A(T48K):C also showed activity similar to wild-type, which we attribute to the ability of the long lysine side chain to project up to make favorable interactions with the electronegative lipid glycerol backbone. In contrast, three other charge mutations (LRRC8A(T48D):C, LRRC8A(V47D):C, and LRRC8A(V47K):C) all increased basal channel activity in isotonic solutions and correspondingly reduced activation by hypotonic swelling (by ∼13, 12, and 3-fold, respectively), similar to the activating effects of C-terminal fusions (Fig. 4b,c and Supplementary Fig. 9). This is consistent our structure-based prediction that electrostatic disruption of lipid binding would relieve channel block and increase basal activity.

## Discussion

Our data support a model for LRRC8 channel gating in which lipid embedded within the pore blocks conduction under resting conditions (Fig. 4d). We hypothesize opening an ion pathway through the channel could involve either of two mechanisms (Fig. 4e). In the first possibility, lipids remain associated with the inside of the pore, but channel expansion around the top of the transmembrane helices opens the space between lipid glycerol backbones to permit ion passage. This is unlikely to require large conformational changes, as a symmetric displacement of each subunit and associated pore lipids ∼0.5-1 Å could create enough space for ion passage between lipid glycerol backbones. In the second possibility, pore lipids retreat from inside the channel to the surrounding membrane. This would likely require expansion of intersubunit gaps to permit lipid egress. Such movements are suggested by increased conformational heterogeneity of LRRC8A:C nanodisc structures. It remains to be determined whether disordered N-termini of LRRC8A or LRRC8C subunits contribute directly to gating as suggested by mutational studies^34^. In homomeric LRRC8D channels, ordered N-termini line the walls of the channel and reach to just below the position of pore lipid acyl chains in our LRRC8A:C structures^29^.

The gating model predicts that subunit displacement from the conduction axis would promote channel opening. This is consistent with evidence correlating LRR separation with activity from recordings of channels with bulky C-terminal fusions, which are sterically incompatible with tight LRR association (Fig. 4b,c)^6,31-33^, FRET measurements^33^, and LRRC8A-activating nanobody structures^30^. Intriguingly, heteromeric channels structures presented here display increased LRR separation and subunit displacement compared to homomeric channel structures determined under similar conditions. The unique LRRC8A:C LRR interaction disrupts the LRR interaction network of LRRC8A homomeric channels, necessitates the entire LRRC8C subunit bow outwards, and opens a gap between subunits through the membrane upper leaflet. This may poise heteromeric LRRC8 channels for opening and explain why they display higher channel activity and larger single channel conductance compared to homomeric channels^3,4,24^. Subunit displacement would also likely expand channel cross sectional area. Since area expansion is energetically favored by increased membrane tension, this could explain why activation is promoted by mechanical force^44,45^, for example from fluid injection into cells, membrane blebbing, or hypotonic swelling^6,46-48^.

Intriguingly, structural evidence has recently been reported for lipids bound inside other large pore channels to which LRRC8s are structurally related^49,50^. Lipids from the membrane outer leaflet are observed in cryo-EM structures of pannexin-1, innexin-6, CALHM2, CALHM4, and CALHM6 and have been proposed to be involved in channel gating^51-54^. LRRC8A:C structures presented here suggest embedded lipids could be a common means to block conduction in closed states of large pore channels.

## Methods

### Construct design and protein expression

The sequence for apocytochrome b_562_RIL^37,55^ was codon optimized for *Spodoptera frugiperda* and synthesized (IDT, Newark, NJ). BRIL was then engineered into the extracellular loop between residues 76 and 91 in the *Mus musculus* (mm)LRRC8A vector we used previously for homohexamer structure determination^28^. This generated a construct for expression of mmLRRC8A(76-BRIL-91)- SNS- LEVLFQGP- SRGGSGAAAGSGSGS- sfGFP-GSS-7xHis. The coding sequence for LRRC8C from *Mus musculus* was codon optimized for *Spodoptera frugiperda* and synthesized (Gen9, Cambridge, MA). The sequence was then cloned into a custom vector based on the pACEBAC1 backbone (MultiBac; Geneva Biotech, Geneva, Switzerland) with an added C-terminal TEV protease cleavage site, linker sequence, and mCherry tag, generating a construct for expression of mmLRRC8C-SNS- ENLYFQG- SRGSGSGS- mCherry. These LRRC8A and LRRC8C constructs were iteratively cloned using the I-CeuI and BstXI sites in the pACEBAC1 backbones to generate the dual LRRC8A/LRRC8C expression plasmid. MultiBac cells were then used to generate a Bacmid according to the manufacturer’s instructions. Sf9 cells were cultured in ESF 921 medium (Expression Systems, Davis, CA) and P1 virus was generated from cells transfected with Escort IV reagent (MillaporeSigma, Burlington, MA) according to manufacturer’s instructions. P2 virus was then generated by infecting cells at 2 million cells/mL with P1 virus at a MOI ∼0.1, with infection monitored by fluorescence and harvested at 72 hours. P3 virus was generated in a similar manner to expand the viral stock. The P3 viral stock was then used to infect Sf9 cells at 4 million cells/mL at a MOI ∼2–5. At 72 hours, infected cells containing expressed LRRC8A/LRRC8C proteins were harvested by centrifugation at 2500 x g for 10 minutes and frozen at -80°C.

For electrophysiology, untagged versions of mmLRRC8A and mmLRRC8C were cloned into CMV-promoter IRES vectors with EGFP or mCherry expression reporters respectively. Point mutants were introduced into LRRC8A by PCR. For the tagged version of the complex, mmLRRC8A was cloned to a vector with a CMV-promoter and C-terminal EGFP (mmLRRC8A-SNS-LEVLFQGP-TAAAA-(no start methionine)EGFP-SGGG-10xHis) and mmLRRC8C was cloned into a vector with a CMV-promoter and C-terminal mCherry (mmLRRC8C-SNS-LEVLFQGP-TAAAA-mCherry-SGGG-10xHis).

For single molecule subunit counting, CMV-promoter vectors were made to express the HVCN1 control, mmLRRC8A, and mmLRRC8C tagged with a C-terminal EGFP and HA-tag (i.e., LRRC8A-SNS-LEVLFQGP-TAAAA-(no start methionine)EGFP-SGGG-YPYDVPDYA). For each LRRC8 construct we also made CMV-promoter vectors to express each gene with the same TEV-mCherry tag used for protein purification.

### Channel purification

For purification of LRRC8A/LRRC8C complexes in GDN detergent, cells from 2 L of culture (∼40mL of cell pellet) were thawed in 100 mL of Lysis Buffer containing (in mM) 50 HEPES, 150 KCl, 1 EDTA pH 7.4. Protease inhibitors (Final Concentrations: E64 (1 µM), Pepstatin A (1 µg/mL), Soy Trypsin Inhibitor (10 µg/mL), Benzamidine (1 mM), Aprotinin (1 µg/mL), Leupeptin (1 µg/mL), AEBSF (1mM), and PMSF (1 mM)) were added to the lysis buffer immediately before use. Benzonase (4 µl) was added after cell thaw. Cells were then lysed by sonication and centrifuged at 150,000 x g for 1 hr. The supernatant was discarded, and residual nucleic acid was removed from the top of the membrane pellet using DPBS. Membrane pellets were scooped into a dounce homogenizer containing GDN Extraction Buffer (50 mM HEPES, 150 mM KCl, 1 mM EDTA, 1% Glyco-diosgenin (GDN: Anatrace, Maumee, OH), and all lysis buffer protease inhibitors except PMSF. Membrane pellets were then homogenized in Extraction Buffer (150 mL) and this mixture was gently stirred at 4°C for 2 hrs. The extraction mixture was centrifuged at 33,000 x g for 45 min and the supernatant, containing solubilized membrane protein, was bound to 5 mL of sepharose resin coupled to anti-mCherry nanobody for 1 hr at 4°C. The resin was then collected in a column and washed with 10 mL of Buffer 1 (20 mM HEPES, 150 mM KCl, 1 mM EDTA, 0.01% GDN, pH 7.4), 40 mL of Buffer 2 (20 mM HEPES, 500 mM KCl, 1 mM EDTA, 0.01% GDN, pH 7.4), and 10 mL of Buffer 1. The resin was then resuspended in ∼6 mL of Buffer 1 with 1.5 mM dithiothreitol (DTT) and 1 mg of TEV protease and rotated gently in the capped column overnight at 4°C. Cleaved LRRC8 complexes were eluted with an additional ∼7 mL of Wash Buffer. Eluate was then applied to a column containing 5 mL of sepharose resin coupled to anti-GFP nanobody for a total of five passes through the resin. The resin was then washed with 20 mL of Buffer 1, 10 mL of Buffer 2, and another 10 mL of Buffer 1. The resin was then resuspended in 6 mL of Buffer 1 with 0.5 mg of PreScission protease and rotated gently in the capped column for 2 hrs at 4°C. Cleaved LRRC8A/C heteromeric complexes were then eluted with an additional ∼7 mL of Wash Buffer, spin concentrated to ∼500 µl with Amicon Ultra spin concentrator 100 kDa cutoff (Millipore), and then loaded onto a Superose 6 Increase column (GE Healthcare, Chicago, IL) on an NGC system (Bio-Rad, Hercules, CA) equilibrated in Buffer 1. Peak fractions containing LRRC8A/LRRC8C complexes were then collected and spin concentrated for BRIL antibody binding.

For channel purification and reconstitution into lipid nanodiscs, the initial purification for nanodisc incorporation was performed analogously to the GDN prep with the following modifications: Extraction buffer contained 1% n-Dodecyl-β-D-Maltopyranoside (DDM, Anatrace, Maumee, OH), 0.2% Cholesterol Hemisuccinate Tris Salt (CHS, Anatrace) instead of GDN and extraction time was 3 hrs. Buffers 1 and 2 contained 0.025% DDM and 0.005% CHS in place of GDN. The Superose 6 Increase column buffer contained 0.025% DDM in place of GDN.

Freshly purified complex in column buffer was reconstituted into MSP2N2 nanodiscs with a mixture of lipids and cholesterol (DOPE:POPS:POPC at a 2:1:1 mass ratio with cholesterol added to a molar percentage of 30%, Avanti, Alabaster, Alabama) at a final molar ratio of 1:2:200:86 (Monomer Ratio: LRRC8 complex, MSP2N2, lipid, cholesterol). First, 20 mM solubilized lipid in Nanodisc Formation Buffer (20 mM HEPES, 150 mM KCl, 1 mM EDTA pH 7.4) was mixed with additional DDM detergent and LRRC8 protein. This solution was mixed at 4°C for 30 minutes before addition of purified MSP2N2. This addition brought the final concentrations to approximately 5 µM LRRC8, 10 µM MSP2N2, 1 mM lipid mix with 0.43 mM cholesterol, and 2.35 mM DDM in Nanodisc Formation Buffer. The solution with MSP2N2 was mixed at 4°C for 10 minutes before addition of 48 mg of Biobeads SM2. Biobeads (washed into methanol, water, and then Nanodisc Formation Buffer) were weighed after liquid was removed by pipetting (damp weight). This mix was incubated at 4°C for 30 minutes before addition of another 48 mg of Biobeads (for a total 96 mg of Biobeads per 0.5 mL reaction). This final mixture was then gently tumbled at 4°C overnight (∼ 12 hours). Supernatant was cleared of beads by letting large beads settle and carefully removing liquid with a pipette. Sample was spun for 10 minutes at 21,000 x g before loading onto a Superose 6 Increase column in Nanodisc Buffer (20 mM HEPES, 150 mM KCl, pH 7.4). Peak fractions corresponding to LRRC8 protein in MSP2N2 were collected, 100 kDa cutoff spin-concentrated and used for BRIL antibody binding. MSP2N2 was prepared as described^56^ and His-tags were cleaved with TEV protease.

### BRIL antibody purification and binding

The anti-BRIL Fab BAG2 was generated and purified as described previously^39^. Briefly, *E. coli* BL21 (Gold) cells was transformed with sequence-verified clones of BAG2 in pRH2.2. BAG2 was grown in 2YT media with 100 μg/mL ampicillin at 37°C for 2-2.5 h during which OD600 reached 0.6–0.8, induced with 1 mM IPTG and further grown for 4.5h at 37°C. BAG2 were purified using protein A followed by ion-exchange chromatography. Harvested cells were resuspended in PBS, supplemented with 1 mM PMSF, 1 μg/mL DNase I. The suspension was lysed by ultrasonication. The cell lysate was incubated at 60°C for 30 min. Heat-treated lysate was then cleared by centrifugation, filtered through 0.22 μm filter and loaded onto a HiTrap MabSelect SuRe (GE Healthcare) column pre-equilibrated with 20 mM Tris; pH 7.5, 500 mM NaCl. The column was washed extensively with 20 mM Tris; pH 7.5, 500 mM NaCl followed by elution of the protein with 0.1 M acetic acid. The eluted protein was directly loaded onto an ion-exchange Resource S 1-ml column pre-equilibrated with 50 mM NaOAc; pH 5.0. Column was washed with the equilibration buffer and the bound protein was eluted with a linear gradient 0–50% of 50 mM NaOAc; pH 5.0, 2 M NaCl. Purified BAG2 was dialyzed overnight against 20 mM HEPES; pH 7.5, 150 mM NaCl and its purity analyzed by SDS-PAGE.

The anti-Fab Nb^41,42^ that binds to the hinge region of the light chain of BAG2 was cloned in pET26b+ vector, expressed and purified as described in^38^.

Antibodies were mixed with Buffer 1 and added in molar excess to LRRC8A/LRRC8C complexes for a ratio of approximately 1:3:6 (LRRC8, BAG2, nanobody) and allowed to bind at 4°C for 30 minutes (The ratio of BAG2 to LRRC8A was calculated to be 3-fold or greater in each experiment with the BAG2 to nanobody ratio ∼1:2). The mixture was pelleted at 21,000 x g for 5 minutes before the supernatant was again applied to the Superose 6 Increase column into either Buffer 1 (150 mM KCl), Buffer 1 with 75 mM KCl (low salt), or Nanodisc Buffer to separate bound complex from free antibody. Peak fractions corresponding to the antibody bound complex were then pooled and spin concentrated for grid freezing.

### Cryo-EM sample preparation

Samples were frozen as soon as possible after spin-concentration and a final 21,000 x g, 5-minute spindown to remove any debris. To prepare cryo-EM grids, 2 µl (GDN 150 mM: 2 mg/ml, GDN 75 mM: 1.2 mg/ml) or 3 μL (Nanodisc, 0.7 mg/ml) of sample was applied to a freshly glow-discharged (PELCO easiGlow, Settings: 0.39 mBar, 25 mA, for 25 s) grid (For GDN 150mM: UltrAuFoil 300 mesh R 1.2/1.3, other samples: gold holey carbon Quantifoil 300 mesh R 1.2/1.3). After a ∼5 s manual wait time, the grid was blotted for 3 s (Whatman #1 filter paper) and immediately plunge-frozen in liquid-nitrogen-cooled liquid ethane using a Vitrobot Mark IV (FEI) operated at 4°C and 100% humidity. Grids were clipped after freezing. For grid preparation, the operator was wearing a mask.

### Cryo-EM data collection

All datasets were collected on a Titan Krios G3i electron microscope (Thermo Fisher), operated at 300 kV and equipped with a Gatan BioQuantum Imaging Filter with a slit width of either 15 eV (GDN 150 mM) or 20 eV. Dose-fractionated images (∼50 electrons per Å^2^ applied over 50 frames) were recorded on a K3 direct electron detector (Gatan) using correlated double sampling (CDS) and super-resolution modes with a pixel size of 0.524 Å (1.048 Å physical pixel size). Nine videos (or 18 using 2-shots per hole for the nanodisc sample) were collected around a central hole position using image shift and the defocus was varied from −0.5 to −1.7 µm through SerialEM^57^.

### Cryo-EM data processing

For the 150 mM KCl in GDN sample, 8262 movie stacks were collected, motion-corrected, and binned to 1.048 Å/pixel using MotionCor2 in RELION3.1^58,59^, and CTF-corrected using Ctffind 4.1.13^60^ (Also see Supplementary Fig. 3-6 for additional processing details). Micrographs with a Ctffind reported resolution estimate worse than 6 Å were discarded leaving 6228 for further processing. An initial particle set of 57,893 particles was generated by manual and template-based picking in RELION3.1 and cleaning using the 2D classification, Ab Initio, and Heterogeneous Refinement jobs in cryoSPARC v3.2^61,62^, with all 3D jobs using C1 symmetry. These particles were then used to train Topaz^63^ to pick a set of 723,383 particles. These particles were cleaned using cryoSPARC Heterogeneous Refinements to arrive at a set of 220,907 particles, which were then Non-Uniform (NU) Refined in cryoSPARC, then transferred, refined, and Bayesian Polished in RELION^64^. UCSF pyem tools were used to convert data from cryoSPARC to RELION format^65^. The 75 mM sample was processed similarly to this point with 7208 initial movies pruned to 6228 by eliminating movies with a resolution estimate worse than 6 Å and by manual inspection. Topaz picking was trained with a set of 8,222 particles yielding a dataset of 451,852 particles which was then cleaned, refined, and polished as described previously to yield 216,267 polished particles. For both datasets, subsequent NU-Refinements with polished particles yielded similar maps with relative stability and good resolution throughout the linker, TM, and ECD of the complex, but with heterogeneity in the LRRs and weak BRIL-Fab-nb density marking the LRRC8A subunits.

Heterogeneous refinements in cryoSPARC and Class3D jobs in RELION did not classify and improve LRR heterogeneity. However, 3D Variability and Display jobs in cryoSPARC were able to distinguish two conformations of the complex. By using maps from the 3D Variability Display output as references in Heterogeneous Refinement, we were able to pull apart these two classes (present at a roughly 50:50 ratio in each dataset) and significantly improve the LRR density. At this point, as we were unable to distinguish any major map difference between salt concentrations, so we decided to pool the polished particles, use Heterogeneous Refinement to again pull apart the two conformations, and perform NU Refinement on each. As a final cleaning step, we imported these Refinements into RELION and performed a Class3D job (Three classes, 10 Å initial low-pass filter, no angular sampling, with a Tau of 40). Selection of the higher resolution classes and subsequent NU Refinement generated the final full-particle maps for each conformation. Masked Local Refinements in cryoSPARC generated higher resolution maps for the upper portion of the complex (ECD, TMD, linker) and sets of LRR domains. The particle box size was 416 pixels (binned 2 or 4-fold for initial particle clean-up).

For the Nanodisc sample, processing was similar to the GDN datasets with 3906 initial movies were pruned to 3593 by eliminating movies with a resolution estimate worse than 4 Å. Initial picking was performed in RELION by manual, template-based, and Laplacian-of-Gaussian picking. Topaz picking was trained with a set of 45,582 particles yielding a dataset of 503,916 particles which was then cleaned, refined, and polished as described previously to yield 192,578 polished particles. After a round of Heterogeneous Refinement, the final set of 176,166 particles was NU Refined to yield the final full particle map. Masked Local Refinement was then performed to improve density of the upper portion of the complex.

### Structure modeling, refinement, and analysis

Assignment of LRRC8A and LRRC8C subunits was made using positions of inserted BRIL domains in complex with anti-BRIL Fab BAG2 and anti-Fab nanobody in unmasked maps and confirmed by high resolution features during modeling. Modeling in Coot^66^ and real space refinement was performed in focused refinement maps from cryoSPARC that showed improved density features relative to global refinement maps. Extracellular regions, transmembrane regions, linker regions, waters, and lipids were modeled in top-focused refinement maps that masked out LRR domains (PDBs 8DRK, 8DR8, 8DS9). Cytosolic LRR domains were modeled in LRR-focused refinement maps. For GDN conformation 1 (round LRRs), three LRR-focused maps were used to model five LRR domains: (1) focused on the LRRC8A chain A LRR domain and LRRC8C chain F LRR domain (PDB 8DRN), (2) focused on the LRRC8A chains B and C LRR domains (PDB 8DRO), and (3) focused on the LRRC8A chain C LRR domain (PDB 8DRQ). For GDN conformation 2 (oblong LRRs), one map focused on the LRRC8A chains B and C and LRRC8C chain F LRR domains was used to model three LRR domains (PDB 8DRA). Additional focused refinements on LRR domains did not yield maps with sufficient density features for unambiguous modeling. Models were refined in real space using Phenix^67^ and validated for stereochemical and geometrical suitability with Molprobity^68^. Following refinement in focused maps, global models were generated by rigid body-docking into global refinement maps, merging overlapping elements, and performing B-factor only refinement (PDBs 8DS3, 8DRE, and 8DS9). For display purposes only, composite focused maps were generated by maximum projection in ChimeraX^69^.

The final models consist of 3888, 3069, and 1418 amino acids for GDN conformation 1, GDN conformation 2, and nanodisc structures, respectively. Unmodeled regions include the N-termini (residues 1-14 for LRRC8A subunits and 1-15 for LRRC8C subunits), extracellular loop 1s (residues 69- 91 in LRRC8A subunits, 61-92 in LRRC8C subunits, and the inserted BRIL-Fab-Nb unit in LRRC8A subunits), and intracellular loops (residues 176-229 in LRRC8A subunits and 180-231 in LRRC8C subunits) in all structures. In GDN conformation 1, the LRR domain from chain D is not modeled (residues 409 to C-terminus). In GDN conformation 2, the LRR domains from chains C, D, and E are not modeled. In the nanodisc structure, all LRR domains are unmodeled and linker regions from chains A,B,E, and F are unmodeled.

Cavity and tunnel measurements were made with HOLE^70^. Figures were prepared using ChimeraX, Prism, Adobe Photoshop, and Adobe Illustrator software.

### Electrophysiology

HeLa LRRC8ABCDE -/- cells^26^ and were cultured in DMEM (Gibco, Thermo Fisher Scientific) with 10% FBS and 100 units/mL penicillin and 100 μg/mL streptomycin. Trypsinized cells were deposited on 12 mm glass coverslips in a 6-well dish 1 day prior to transfection. For transfection, LRRC8A and LRRC8C plasmids were mixed with Fugene 6 in OptiMEM at 1ug/well/construct (or 0.5 ug for C-terminally tagged constructs) and applied to cells in growth media without antibiotics. Control cells were not transfected. After 24 hours the transfection media was replaced with growth media with antibiotic and patching was conducted from 26-60 hours post-transfection. The optimal time for achieving whole cells, particularly for constructs with increased basal activity, was ∼30-48 hours post-transfection.

For patching, coverslips were placed in a perfusion chamber at room temperature in isotonic bath solution: in mM, 90 NaCl, 2 KCl, 1 CaCl_2_, 1 MgCl_2_, 10 HEPES (NaOH), 10 Glucose, ∼130 Mannitol adjusted to a final pH 7.4 with NaOH. Mannitol was used to adjust the solution osmolarity to ∼330 mOsm, the approximate osmolarity of the cell culture media. Solution osmolarity was measured with a vapor pressure osmometer (Vapro #5600, ELITechGroup) and buffers are roughly adapted from^26^. Cells were chosen by a combination of the presence of the fluorescent reporters or tags (both GFP and mCherry) and cell morphology (aiming for healthy interphase cells). Borosilicate glass pipettes were pulled to a resistance of ∼2.5-3 MΩ and filled with pipet solution: in mM, 133 CsCl, 5 EGTA, 2 CaCl_2_, 1MgCl_2_, 10 HEPES, 4 MgATP, adjusted to pH 7.3 with CsOH (solution measured at ∼330 mOsm). An Axopatch 200B amplifier connected to a Digidata 1440B digitizer (Molecular Devices) was used for data acquisition with pClamp10.7 software. Analog signals were filtered at 1 kHz and sampled at 100 kHz. Pressure application from the patch pipette was accomplished with a high-speed pressure clamp (HSPC, ALA Scientific). Once a whole-cell patch clamp mode was achieved and stable whole-cell capacitance was measured, voltage families were recorded to monitor pre-swell currents with the following voltage protocol applied once every 2 s: V_hold_ = 10 mV; V_test_ = −100 to +100 mV, Δ20 mV, 250 ms. When initial currents stabilized, hypotonic bath solution (same components as isotonic, but adjusted to ∼250 mOsm by addition of ∼40 mM Mannitol) was exchanged into the chamber and currents were monitored over the course of cell swelling until currents stopped increasing for multiple records or the patch broke. If a patch exhibited signs of leaking or sealing, a membrane test protocol was examined to either reseal, re-break-in, or discard the patch. Care was taken to make sure that the measured pressure was slightly negative (i.e., -1 mm Hg) after whole-cell mode was achieved. Cells were monitored on the brightfield microscope and were not used if the cell showed signs of death, swelling, or blebbing prior to exchange into the hypotonic buffer.

Data was analyzed using Clampfit 10.7, Excel, and Graphpad Prism software. For voltage families the data was decimated at 100X and plotted in Prism. Displayed data are current densities from each cell recorded at 60 mV after achieving stable currents in isotonic and hypotonic solutions and the ratio of these current densities (fold activation). Statistical analysis between wild-type and mutant or tagged LRRC8A/C was conducted with Dunnett’s multiple comparisons test after an ordinary one-way ANOVA using Prism software.

### Single molecule subunit counting

Fluorescent protein-tagged constructs for photobleaching were transfected into LRRC8 knockout HEK293 cells at 2µg/construct/well using Fugene 6. For the HVCN1 and LRRC8A controls, we used cells from 1-well of a 6-well plate per preparation and for the LRRC8A:C double transfections we used two wells worth of cells. Two days after transfection, cells were harvested by centrifugation at 1000 x g for 5 minutes and frozen at -80°C. For preparation of protein, cells were resuspended in Lysis Buffer (same as large-scale prep but with 20 mM HEPES) and lysed by sonication. The membrane was centrifuged for 5 minutes and 21,000 x g, then the supernatant was discarded and the pellet rinsed with Lysis Buffer. The membrane pellet was resuspended in GDN Extraction Buffer (same as large scale prep but with 20 mM HEPES) using a dounce homogenizer and rotated at 4°C for 1 hour. After extraction, samples were centrifuged for 5 minutes at 21,000 x g and the supernatant was collected. The control samples were then directly used for subunit counting. For the LRRC8 heteromer constructs, the supernatant was then bound to 50 µl of mCherry nanobody resin by rotation at 4°C for 1 hour. After bead binding, the sample was rinsed twice in GDN Wash Buffer (Buffer 1), then washed twice by rocking at 4°C for 5 minutes. Following the final wash, the beads were resuspended in GDN Wash Buffer and LRRC8 heteromers were eluted by addition of 20 µg of TEV protease with 1 mM DTT. The cleavage reaction was performed for 2 hours at 4°C. After elution, samples were centrifuged for 5 minutes at 21,000 x g and the supernatant, containing LRRC8 heteromers, was collected for subunit counting. Care was taken to avoid light exposure during the mini-purification.

Imaging chambers were prepared using mPEG (Laysan Bio) passivated glass slides and coverslips (VWR) doped with biotin PEG16. Imaging chambers were incubated with NeutrAvidin (40ug/mL) (Thermo Fisher) for 15 minutes followed by biotinylated anti-HA (ab26228) antibody (1:100) 1-2 hours. Samples were diluted in GDN Wash Buffer (1-1000x depending on protein concentration), flowed into imaging chamber and incubated briefly (10 s - 5 min) in order to achieve low density immobilization. Unbound protein was extensively washed out of the imaging chamber using GDN Wash Buffer. Samples were imaged on an objective-based (1.49 NA 60x, Olympus) TIRF microscope with a Photometrics Prime 95B sCMOS camera at a 100 ms frame rate. A 491 nm laser (Cobolt Calypso) was used for EGFP excitation and 500 frames were recorded for each movie to ensure bleaching of each EGFP. Micro-Manager was used for data acquisition. Movies were collected in two separate days of experiments.

Six movies from each day were analyzed, each containing a minimum of 50 fluorescent spots. Fluorescence traces were extracted from raw data files using SPARTAN 3.7.0 single-channel settings and bleaching steps were determined manually. Due to the increased occurrence of near simultaneous bleaching events and resultant difficulty in accurately discriminating step number, traces with over three bleaching steps were combined into a single category. Traces with three or more bleaching steps were combined for HV1 to match previously published data. A clear number of bleaching steps could not be determined at all for approximately 5-10% of traces and these traces were therefore excluded from plotted data. Excel and Graphpad Prism software were used for data analysis and graphing. Binomial distribution predictions were made for LRRC8A-EGFP using varying probabilities of functional GFP assuming that only homohexamers were present. For LRRC8A:C heteromers, binomial predictions were made assuming 60% functional GFP for a fixed 3:3 stoichiometry as well as for all possible combinations. In the latter case, the binomial distribution was calculated with n=6 and p=(.5)(.6) to account for equal likelihood of each subunit as well as GFP functionality. Formation of complexes containing zero and six of each subunit were permitted in initial calculations and distributions were subsequently normalized to exclude these combinations that we could not observe in this assay due to lack of fluorescence and the mini-purification.

## Data Availability

For LRRC8A_BRIL_:C in GDN conformation 1, final models are in the PDB under 8DS3,8DRK,8DRN,8DRO and 8DRQ and final maps are in the Electron Microscopy Data Bank (EMDB) under EMD-27682, 27677, 27677, 27679, and 27681. For LRRC8A_BRIL_:C in GDN conformation 2, final models are in the PDB under 8DRE, 8DR8, and 8DRA and final maps are in the Electron Microscopy Data Bank (EMDB) under EMD- 27676, 27674, and 27675. For LRRC8A_BRIL_:C in lipid nanodiscs, final models are in the PDB under 8DSA and 8DS9 and final maps are in the Electron Microscopy Data Bank (EMDB) under EMD-27687 and 27686. Original micrograph movies and final particle stacks are in the Electron Microscopy Public Image Archive (EMPIAR).

## Acknowledgements

We thank J. Remis, D. Toso, and P. Tobias for microscope and computational support at the Cal-Cryo facility. We thank the labs of A. Patapoutian and T.J. Jentsch for gifts of HeLa and HEK293 LRRC8 knockout cell lines. We thank members of the Brohawn laboratory and A.F. Kern, C.M. Hoel, and R.A. Rietmeijer for discussions and feedback on the manuscript. S.G.B. is a New York Stem Cell Foundation-Robertson Neuroscience Investigator. This work was additionally funded by NIGMS grant GM128263 to D.M.K., a McKnight Foundation Scholar Award, a Sloan Research Fellowship, and a Winkler Family Scholar Award (to S.G.B.).

## Author Contributions Statement

D.M.K. performed and analyzed all biochemistry, cryo-EM, and electrophysiology experiments. J.M.H. contributed to development of fiducial tag constructs. S.M. and A.A.K. provided anti-BRIL Fab and anti-Fab Nb. J.B. performed subunit counting experiments and analyzed data under supervision from E.Y.I. D.M.K. and S.G.B. modeled and refined the structures. D.M.K. and S.G.B. conceived of the project. S.G.B. secured funding and supervised the project. D.M.K. and S.G.B. wrote the manuscript with input from all authors.

---

**Supplementary Figure 1.**
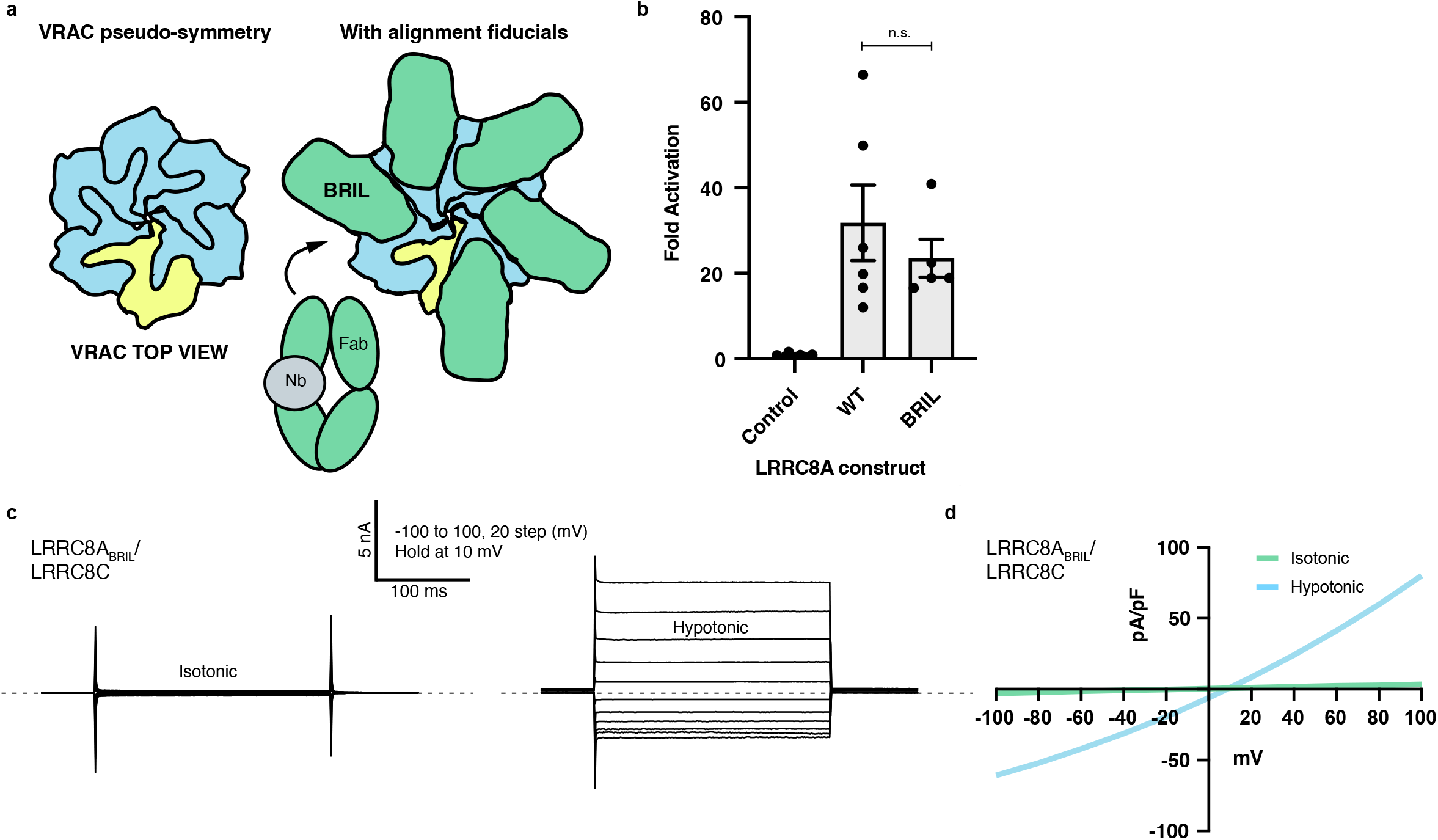
Fiducial tagging strategy. **(a)** Schematic of biochemical approach. A BRIL domain is inserted into the extracellular loop 1 as a fiducial to facilitate computational alignment of pseudosymmetric LRRC8A and LRRC8C subunits in single particle cryo-EM reconstructions. Anti-BRIL Fabs and anti-Fab Nbs are added to purified samples to increase fiducial mass. **(b)** Function of LRRC8A-BRIL:C channels. Comparison of fold activation by hypotonic solution from wild-type LRRC8A:C and LRRC8A-BRIL:C expressing LRRC8A/B/C/D/E -/- cells. No statistically significant difference is observed (Student’s t-test, P=0.45, n=6 and 5 cells for LRRC8A:C and LRRC8A-BRIL:C, respectively). **(c)** Representative voltage clamp recording from LRRC8A-BRIL:C expressing LRRC8A/B/C/D/E -/- cells in isotonic (left) and hypotonic (right) solutions. **(d)** Current-voltage relationship from data in **(c)**.

**Supplementary Figure 2.**
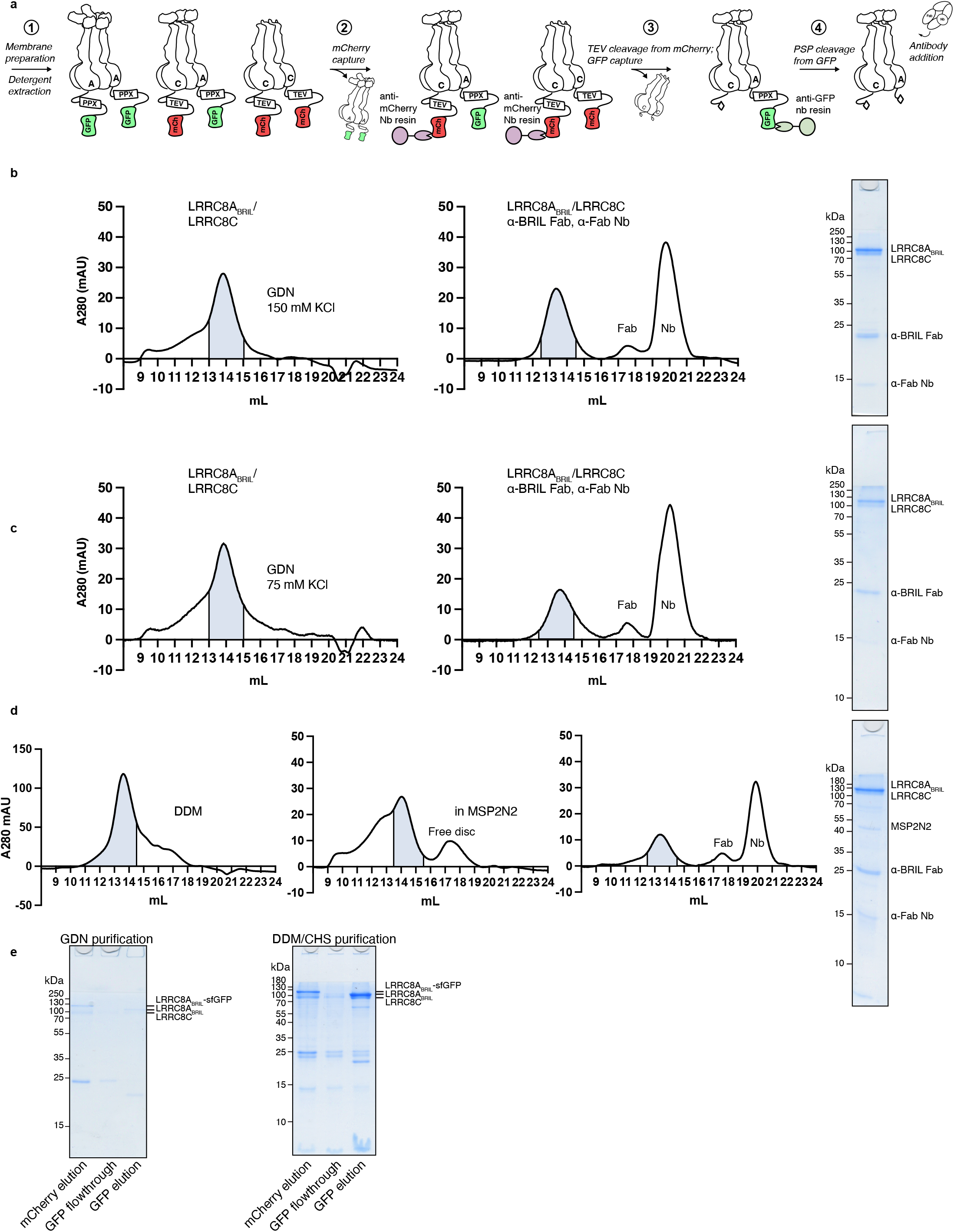
Purification and reconstitution of LRRC8A:C channels. **(a)** Schematic of purification strategy. **(b)** Superose 6 gel filtration chromatogram of LRRC8A_BRIL_:C purified in GDN and 150 mM KCl before (left) and after (middle)) addition of anti-BRIL Fab and anti-Fab Nb with pooled fractions indicated in blue and Coomassie stained SDS–PAGE of final sample (right). **(c)** Superose 6 gel filtration chromatogram of LRRC8A_BRIL_:C purified in GDN and 75 mM KCl before (left) and after (middle)) addition of anti-BRIL Fab and anti-Fab Nb with pooled fractions indicated in blue and Coomassie stained SDS–PAGE of final sample (right). **(d)** Superose 6 gel filtration chromatogram of LRRC8A_BRIL_:C purified in DDM, after reconstitution in MSP2N2 lipid nanodiscs, and after addition of anti-BRIL Fab and anti-Fab Nb with pooled fractions indicated in blue and Coomassie stained SDS–PAGE of final sample (right). **(e)** Coomassie stained SDS–PAGE of samples from purification in GDN (left) and DDM/CHS (right) prior to gel filtration chromatography. Note presence of LRRC8C band in anti-GFP nanobody flowthrough indicating presence of homomeric LRRC8C channels.

**Supplementary Figure 3.**
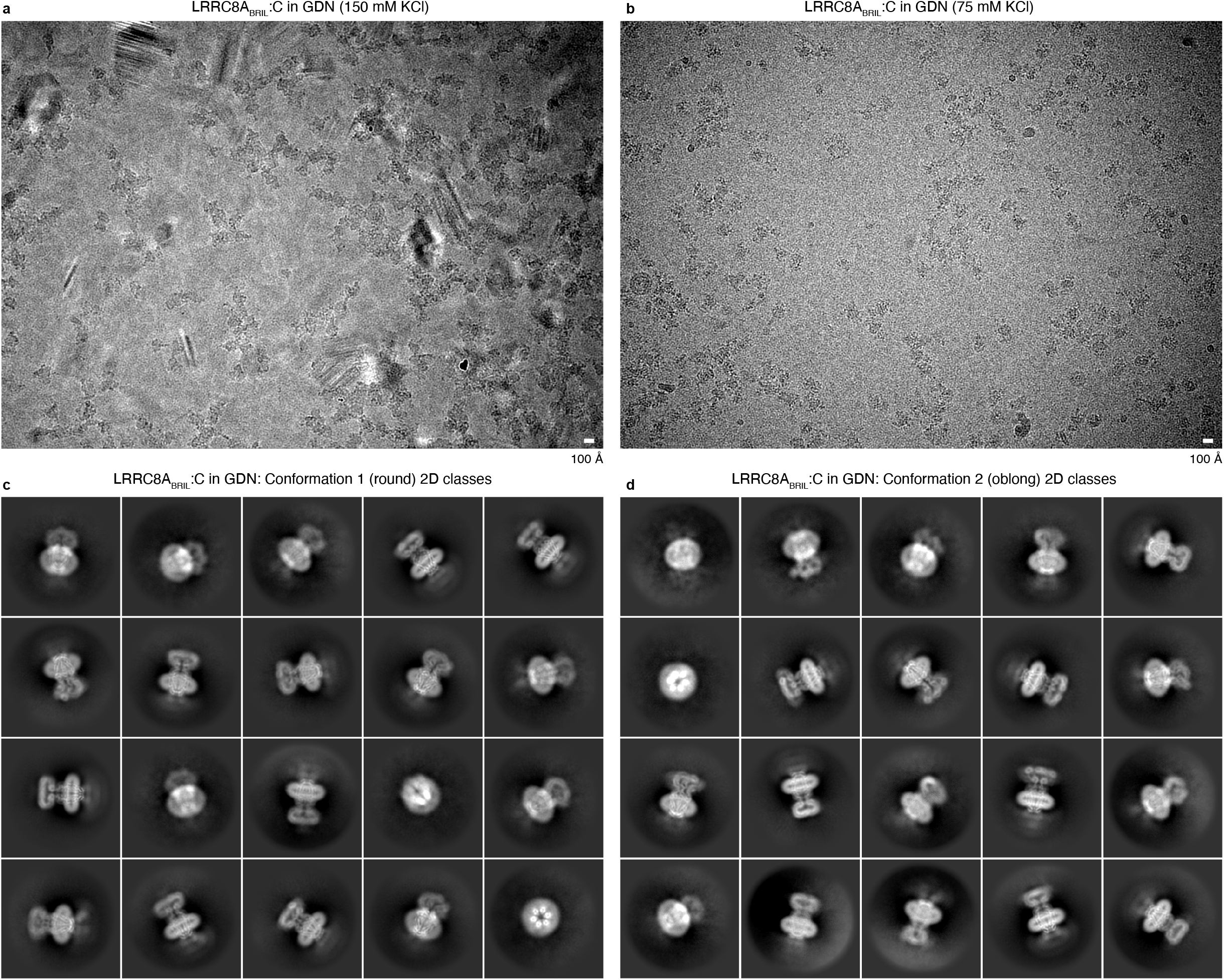
Cryo-EM data for LRRC8A_BRIL_:C in GDN. **(a**,**b)** Representative micrographs from data collected of LRRC8A_BRIL_:C in GDN and **(c**,**d)** selected 2D class averages for conformations 1 and 2.

**Supplementary Figure 4.**
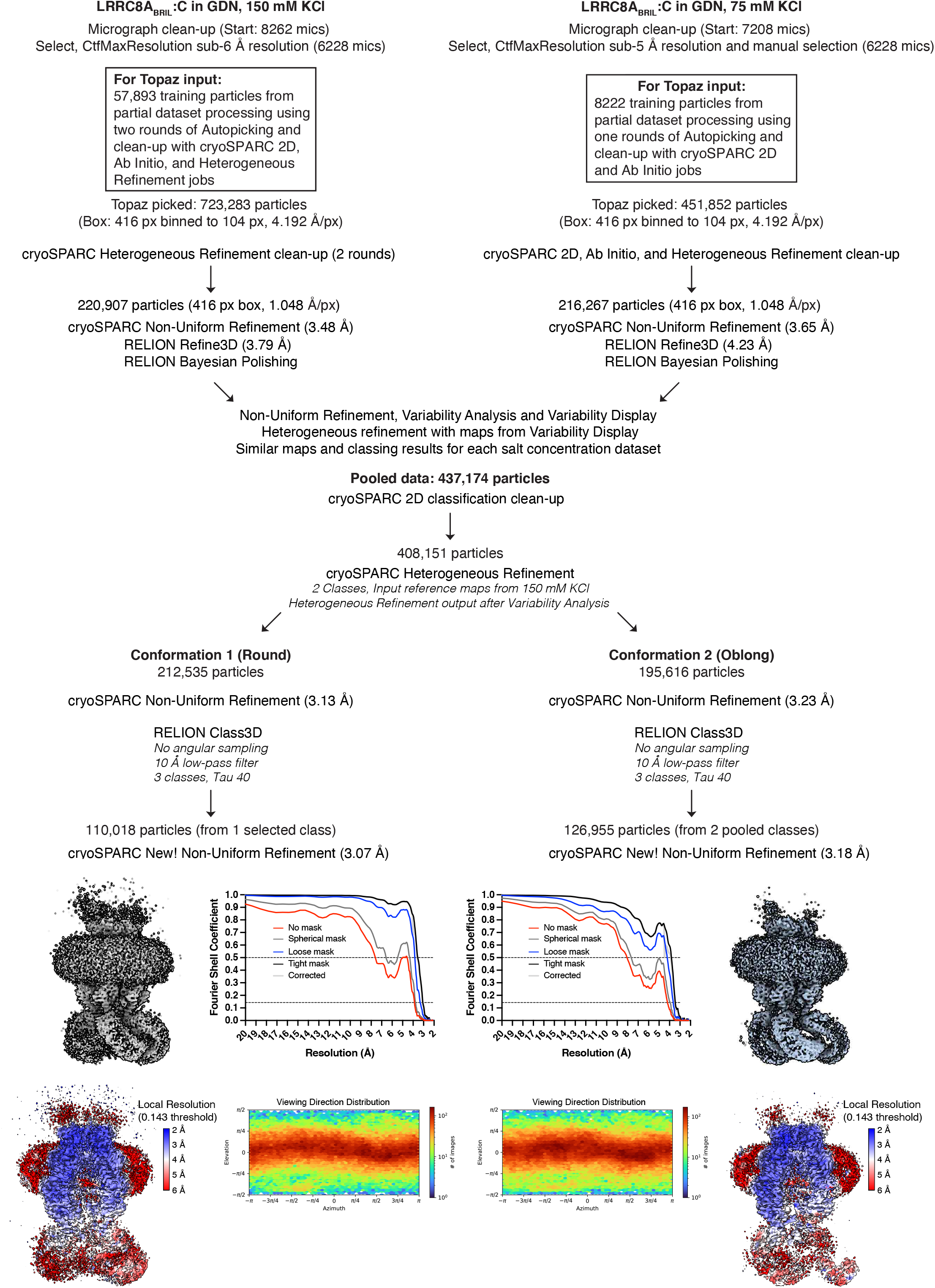
Cryo-EM processing pipeline and validation for LRRC8A_BRIL_:C in GDN. (a,b) Cryo-EM data processing steps for conformation 1 and 2

**Supplementary Figure 5.**
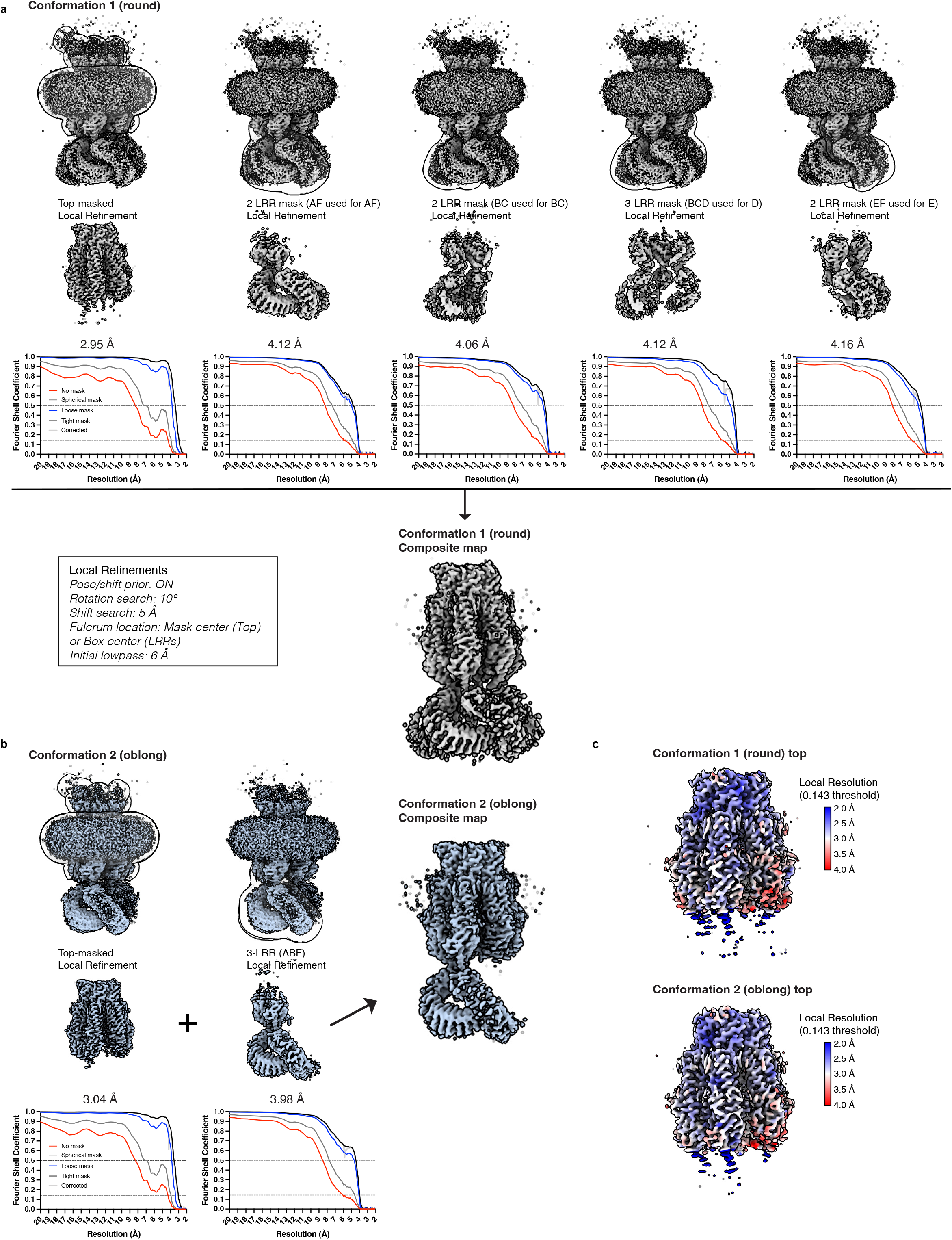
Focused refinements and validation for LRRC8A_BRIL_:C in GDN. (a,b) Focused refinement strategy for conformation 1 and 2.

**Supplementary Figure 6.**
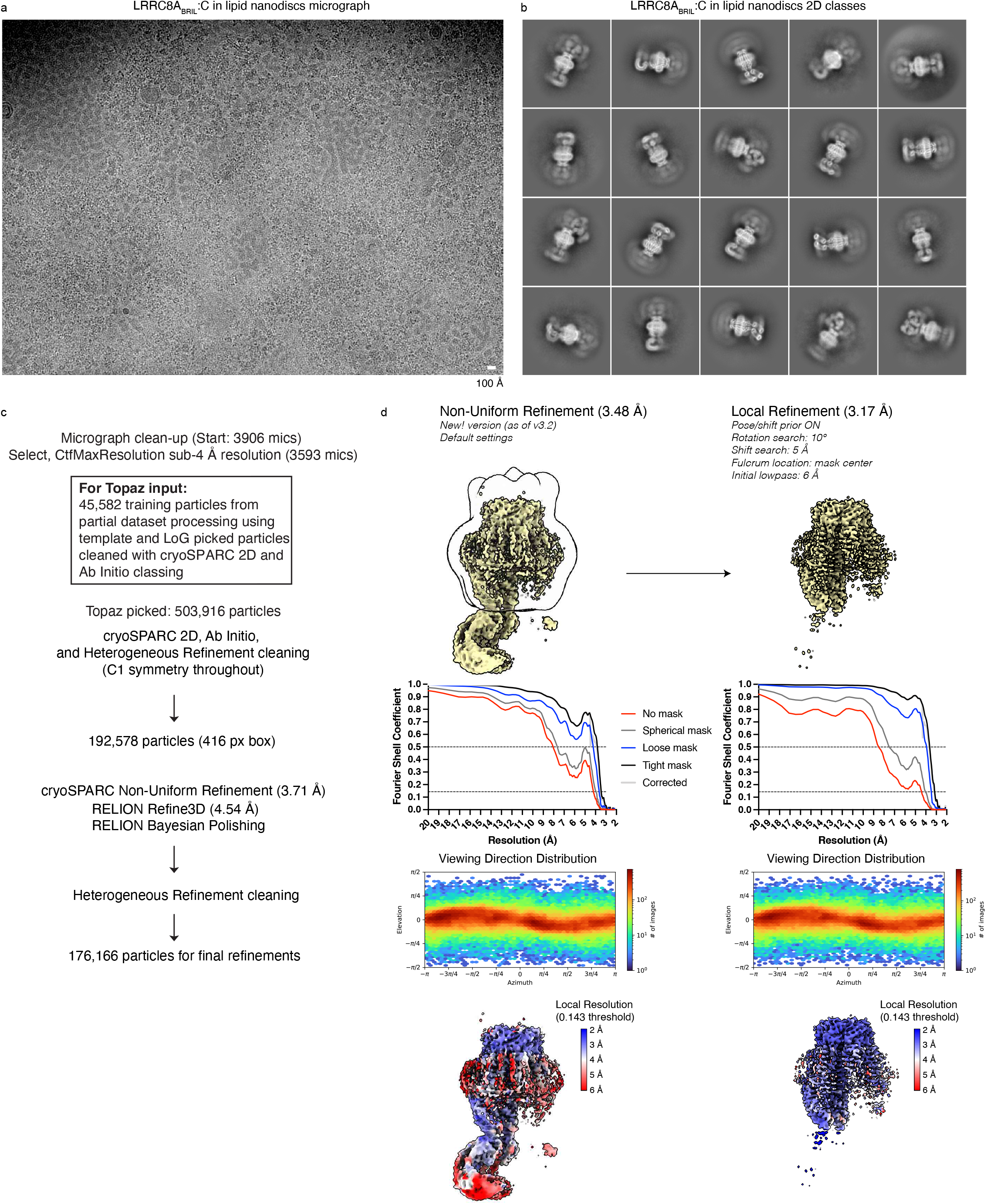
Cryo-EM data and validation for LRRC8A_BRIL_:C in lipid nanodiscs. **(a)** Representative micrograph, **(b)** selected 2D class averages, **(c)** data processing pipeline, and **(d)** validation and focused refinement strategy for LRRC8A_BRIL_:C in lipid nanodiscs.

**Supplementary Figure 7.**
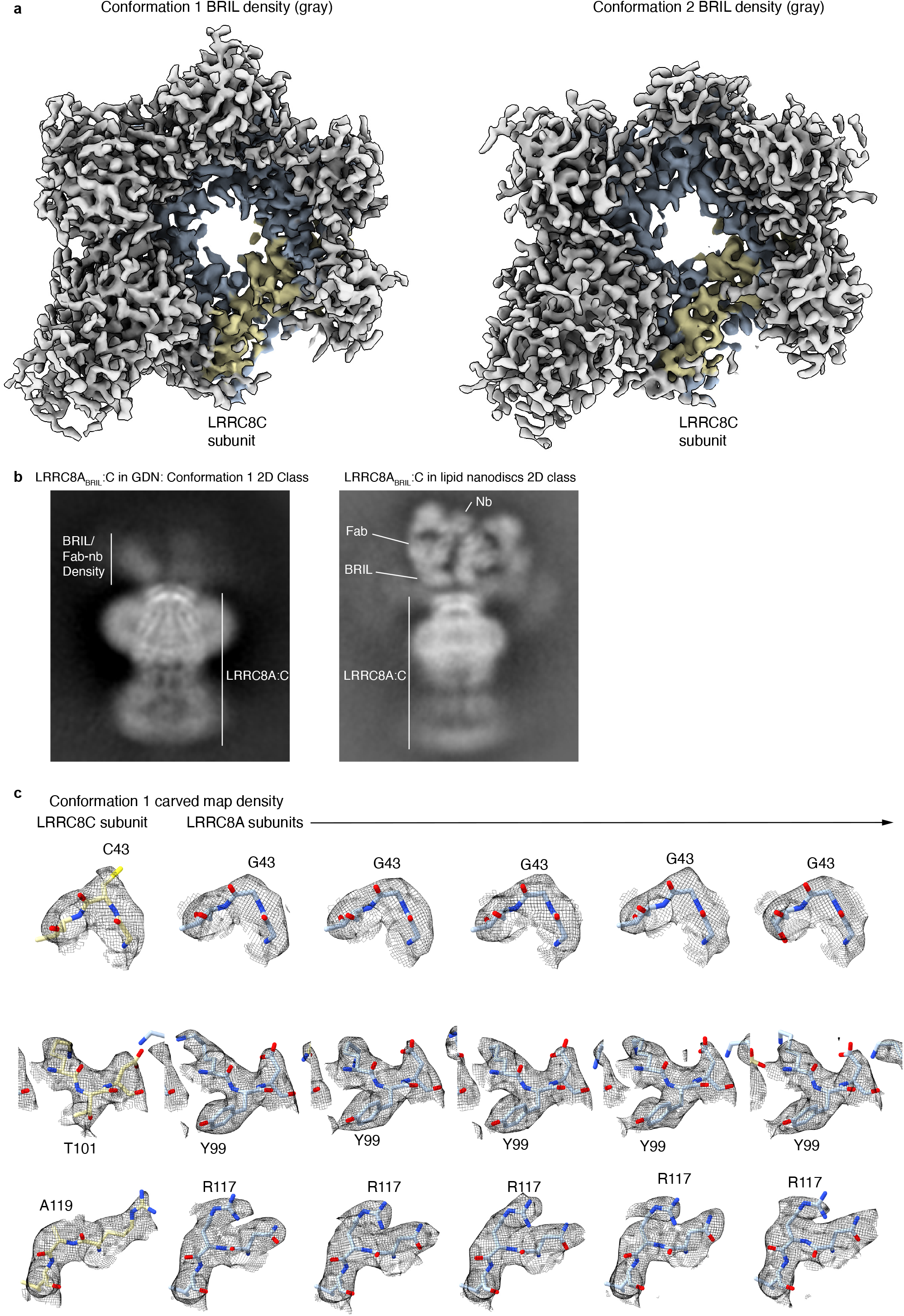
Cryo-EM maps and validation of subunit assignment. **(a)** Views of cryo-EM maps from the extracellular side at low contour illustrating position of density for BRIL-Fab-Nb (gray) above five LRRC8A subunits (blue) and one LRRC8C subunit (yellow). **(b)** Selected 2D Classes with labeled BRIL-Fab-Nb densities. **(c)** Selected regions of cryo-EM density around amino acids that are different in LRRC8A and LRRC8C subunits.

**Supplementary Figure 8.**
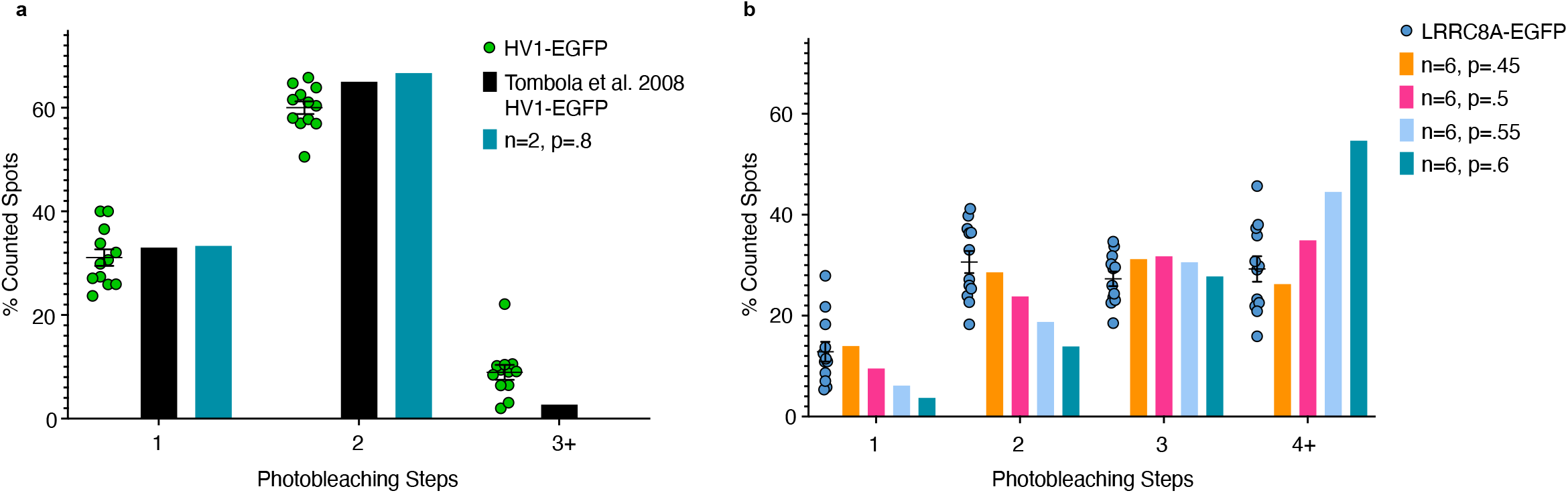
Validation of EGFP subunit counting strategy. **(a)** Distribution of HV1-EGFP photobleaching steps compared to previously published data^43^ and a predicted binomial distribution with 80% functional EGFP. **(b)** Distribution of LRRC8A-EGFP photobleaching steps and predicted binomial distributions for EGFP-tagged homohexamers with varying ratios of mature and fluorescent EGFP (mean ± sem for n=12 movies with ≥ 50 particles each).

**Supplementary Figure 9.**
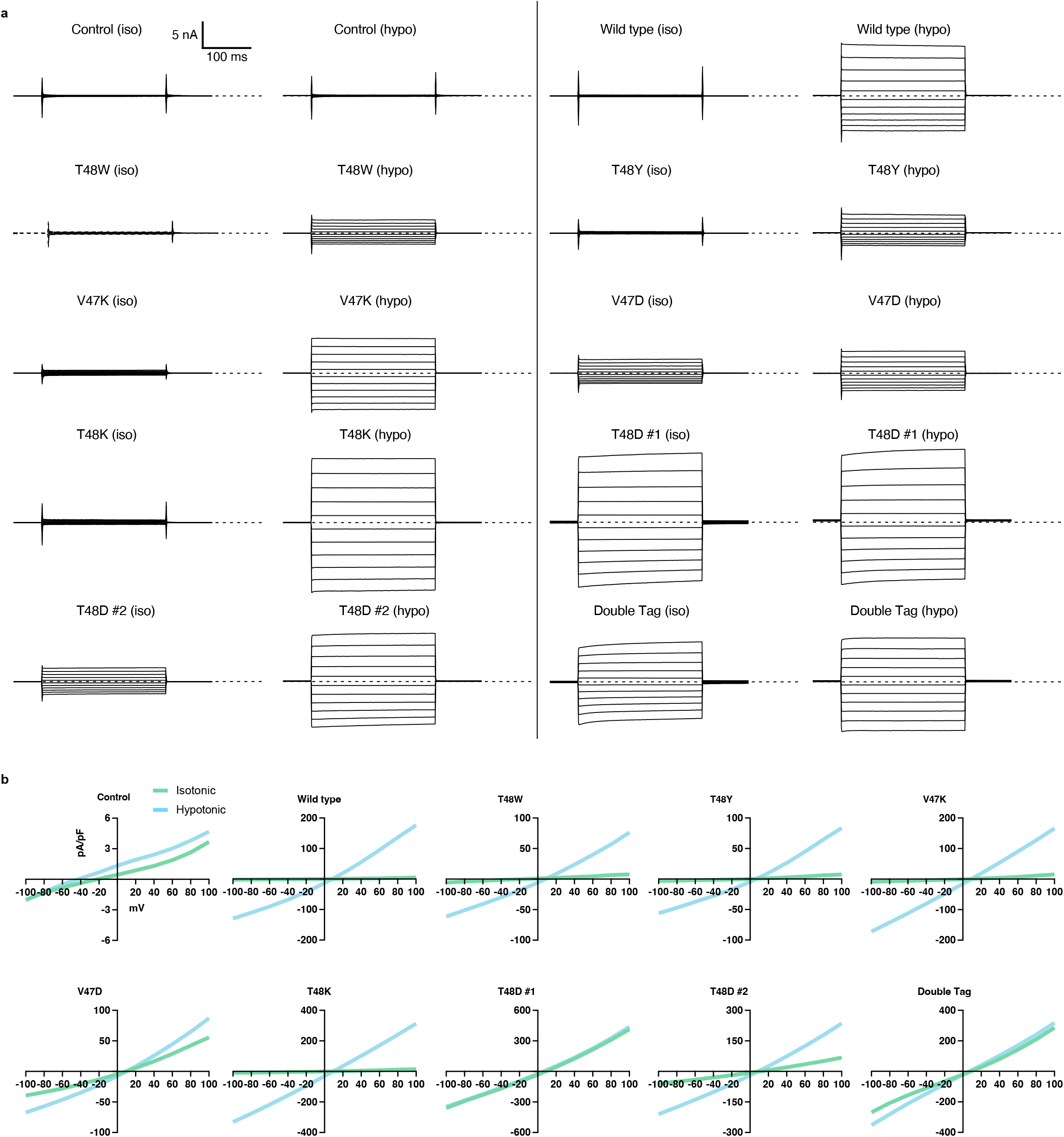
Functional characterization of LRRC8A:C mutants. **(a)** Representative voltage family recording from each construct tested in isotonic and hypotonic solution.**(b)** Current-voltage relationship from data in **(a)**.

